# O-GlcNAcylation drives macrophage IL-4 responsiveness and tissue residency through metabolic and cell cycle calibration

**DOI:** 10.64898/2026.01.05.697622

**Authors:** Graham A. Heieis, Conor M Finlay, Thiago A Patente, Martina Erbì, Thijs van der Meer, Jordan O’Keeffe, Joost M Lambooij, Frank Otto, Frank Faas, Aat A Mulder, Rick Maizels, Roman I Koning, Bart Everts

## Abstract

The metabolic requirements for macrophage IL-4 polarization remain contentious, while immunometabolic studies of tissue resident macrophages are still sparse. Hexosamine biosynthesis has gained attention regarding its immune regulatory potential via downstream O-GlcNAcylation. Here we identify protein O-GlcNAcylation as a requirement for IL-4 polarization *in vitro* and proliferative expansion *in vivo* during cytokine challenge or infection. We further show that O-GlcNAcylation is critical for controlling tissue residency. By enforcing metabolic and cell cycle quiescence during differentiation, O-GlcNAcylation is needed for adult monocytes to establish a long-live residency program. In this context, its absence leads to perpetual DNA vulnerability and damage via reactive oxygen species, resulting in a senescent-like state poised for cell death. Conversely, long-lived populations instead require O-GlcNAcylation for self-renewal and inflammatory expansion. Our findings altogether suggest O-GlcNAcylation, fueled by hexosamine biosynthesis, serves as a central metabolic rheostat for resident macrophage formation and maintenance during homeostasis and disease.

## INTRODUCTION

Cellular metabolism is an integral component of macrophage function. The importance of metabolic regulation in macrophages is clearly illustrated using *in vitro* polarization models of either inflammatory activation with bacterial ligands such as LPS, or alternative activation via the cytokine interleukin-4 (IL-4). In this dichotomy, classical activated (murine) macrophages engage glycolysis^1,2^, the pentose phosphate pathway^3^ and lipid synthesis^4,5^, while mitochondrial oxidative phosphorylation (OxPhos) is nullified, largely by the activity of nitric oxide synthase and associated respiratory burst^2,6^. The metabolic faculties of Il-4 polarized, alternatively activated macrophages (AAM) remain more enigmatic. While AAM increase glycolysis^7^, glutaminolysis^8^, fatty acid oxidation^9,10^, reports are contradictory as to their nutrient requirements for AAM differentiation and function^11–14^.

Recently the hexosamine biosynthesis pathway (HBP) has garnered attention as a regulator of macrophage responses^15^, and is a feature of AAM metabolism relative to classically activated macrophages. The HBP product UDP-GlcNAc can be added to proteins by N-linked glycosylation, and this was initially proposed as a requirement to fully promote AAM polarization^16^. More Recent attention has turned to O-linked glycosylation in the form of O-GlcNAcylation, an intracellular post-translational modification analogous to phosphorylation, involving the addition of GlcNAc residues by the activity of O-GlcNAc transferase (OGT). Some studies suggest O-GlcNAcylation may also be important in AAM function by potentiating STAT6 activity^17,18^, but are met with conflicting reports suggesting AAM are unaffected when O-GlcNAcylation is blocked^19,20^. Further complexity is added by findings that bacterial^21,22^ and viral stimuli^23–25^ also lead to altered flux through the HBP.

Despite elegant molecular findings, studies on macrophage O-GlcNAcylation have relied on *in vitro* systems that lack the complexity of *in vivo* settings, namely the diversity in macrophage ontogeny within tissues. Peripheral organs are seeded with macrophages during early developmental stages of life and maintained throughout adulthood by a combination of local proliferation and replacement by circulating monocytes^26^. The turnover kinetics of macrophages vary according to tissue and microanatomical location; however, resident macrophages are frequently depleted during inflammation and replaced by monocyte-derived macrophages with resolution^27–30^. Studies have pointed towards dynamic metabolic changes during macrophage differentiation^31–33^, yet the exact mechanisms that regulate this reprogramming have largely gone unexplored. The HBP is metabolically demanding, requiring integration of glutamine, glucose, nucleotide and fatty acid pathways, and thus serves as an intrinsic nutrient sensor for the cell^34^. Accordingly, metabolic and cell-cycle proteins are common targets whose activity is regulated by O-GlcNAcylation^34,35^. Thus, there is precedent to more thoroughly interrogate the *in vivo* role of O-GlcNAcylation in tissue macrophages.

Here, we initially investigated the contribution of O-GlcNAcylation to AAM polarization. We found that O-GlcNAcylation was required for optimal IL-4 responses *in vitro* and *in vivo* during cytokine stimulation or helminth infection but also revealed a profound decrease in tissue resident populations. We propose O-GlcNAcylation temporally controls monocyte-derived macrophage differentiation by promoting metabolic and cell-cycle quiescence upon entry into the tissue. In its absence, tissue macrophages acquire an inflammatory, senescent phenotype that is prone to cell death due to metabolic stress. We therefore identify O-GlcNAcylation as a novel regulator of both IL-4-driven alternative activation and tissue residency of macrophages.

## RESULTS

### O-GlcNAcylation promotes macrophage IL-4 responsiveness in vitro and in vivo

The addition of O-GlcNAcylation to proteins is controlled by O-GlcNAc Transferase (OGT), while removal is performed by O-GlcNAcase (OGA) (**Figure 1A**). Pharmacologically blocking OGT with the inhibitor ST045849 (OGTi) suppressed AAM polarization of bone marrow derived macrophages (BMDM) at the protein and transcriptional level, without impairing the response to IFNγ/LPS (**Figure 1B & C**). To definitively address the importance of the O-GlcNAcylation in AAM, we generated mice with a macrophage specific deletion of *Ogt*, by crossing *Lyz2*-*cre* to *Ogt-flox* mice. We then confirmed that BMDM cultured from *Lyz2^ΔOgt^* mice showed a reduction, albeit incomplete, of total O-GlcNAc staining (**Figure 1D**). Nevertheless, we similarly observed *Lyz2^ΔOgt^* BMDM had a marked decrease in AAM polarization (**Figure 1E**). O-GlcNAcylation in *Ogt*-sufficient BMDM peaked within 8 hours of IL-4 stimulation but remained low in the absence of *Ogt* (**Figure 1F**). Accordingly, treatment with OGTi had the strongest inhibitory effect when added within the first 8 hours of stimulation (**Figure 1F**). Despite the significant impairment of AAM polarization, we did not see differences in STAT6 tyrosine phosphorylation (**Figure 1G**). Thus, OGT is critically required for IL-4-driven polarization *in vitro*, without an observable effect on STAT6 activation.

**Figure 1.**
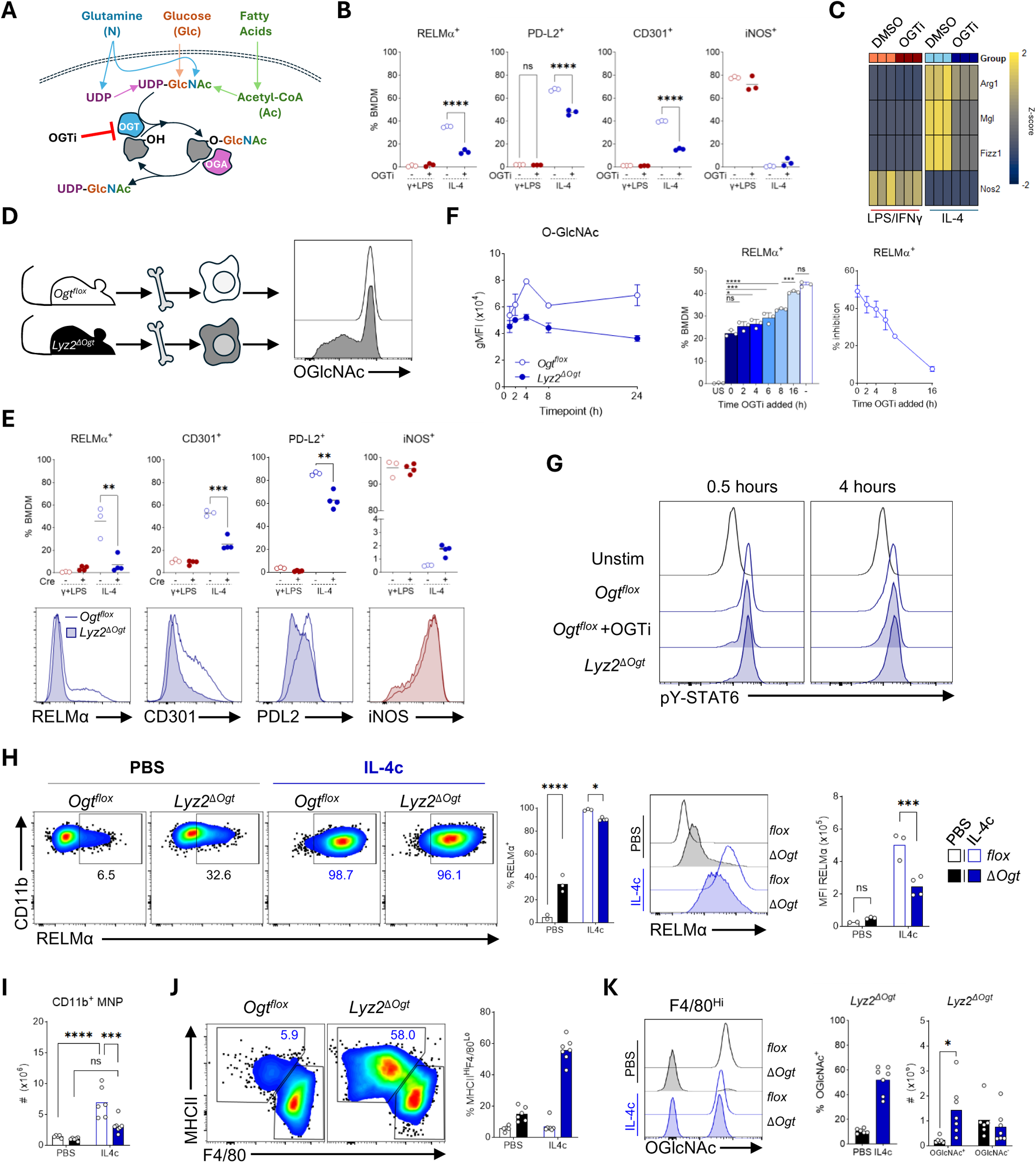
O-GlcNAcylation promotes alternative activation of macrophages. (**A**) Overview of the hexosamine biosynthesis pathway (HBP) and O-GlcNAc cycling between O-GlcNAc transferase (OGT) and O-GlcNAcase (OGA). (**B** and **C**) Stimulated BMDM treated with 20µM of ST045849 (OGTi), assessed by (B) flow cytometry and (C) qPCR. Representative of three experiments, n=3 mice/group. (**D**) Representative detection of O-GlcNAc in d7 differentiated BMDM. (**E**) Expression of activation markers in *Ogt^flox^* or *Lyz2^ΔOgt^* BMDM stimulated for 24 hours. Representative of three experiments, n=3-4 mice/group. (**F**) Time course of O-GlcNAc detection following IL-4 stimulation and RELMα^+^ BMDM following different times of OGTi addition. Representative of two experiments. (**G**) Tyrosine 641 phosphorylation of STAT6. Representative of two experiments. (**H-K**) 5µg of IL-4c injected at d0 and d2 followed by analysis of PEC on d3 for (**H**) RELMα expression, (**I**) number of MNP, (**J**) frequency of MNP populations and (**K**) O-GlcNAc^+^ cells. One of two representative experiments shown (**H**) or pooled (**I-K**), n=2-4 mice/group. Data presented as mean with biological replicates shown. Statistical comparison done with students (B and E) unpaired t-test, (F) one-way ANOVA, or (H-K) two-way ANOVA, with Tukey’s correction.

To test whether OGT also controls AAM formation *in vivo*, we injected IL-4:IgG complex (IL-4c) intraperitoneally and harvested the peritoneal exudate cells (PEC) by lavage. The peritoneal cavity contains multiple populations of mononuclear phagocytes (MNPs)^36^ with two primary macrophage populations: MHCII^Hi^F4/80^Lo^ small cavity (SCM) and MHCII^-^F4/80^Hi^ large cavity macrophages (LCM). In mice, SCM are thought to be largely monocyte-derived, whereas LCM are primarily of embryonic origin but supplanted by monocytes with aging, in part by the progressive differentiation of SCM via a converting intermediate (CCM), which can be expedited by inflammation^29,36^. DC-like macrophages can be identified by CD11c expression, and here likely contains a mix of cDC2 and early macrophage progenitors^36,37^.

LCM from *Lyz2^ΔOgt^* mice expressed significantly less RELMα and failed to expand in response to IL-4c, supporting a block in AAM activation *in vivo* (**Figure 1H & I**). Although there was minimal change in the frequency of RELMα^+^ cells compared to *Ogt^flox^* mice, these observations were confounded by an increase in RELMα expression in PBS treated *Lyz2^ΔOgt^* mice; an indication of recent SCM conversion at steady-state^37,38^ (**Figure 1H**). Accordingly, after IL-4c injection, the MNPs in *Lyz2^ΔOgt^* mice shifted to a dominant composition of MHCII^Hi^ SCM/CCM, consistent with a switch from local proliferation to monocyte recruitment (**Figure 1I**). Additionally, 70% of remaining LCM were O-GlcNAc^+^, pointing towards recent derivation from hematopoietic precursors (**Figure 1K**). Furthermore, the number of O-GlcNAc^-^ cells was unchanged, supporting a failed proliferative response to IL-4 (**Figure 1K**). These data therefore reveal that O-GlcNAcylation is required for complete LCM polarization and proliferation in response to IL-4 *in vivo*, leading to an alternate response of monocyte recruitment.

### Macrophage O-GlcNAcylation supports physiological AAM polarization and T cell activation

We then tested whether the altered response to IL-4 has physiological consequences using the intestinal helminth *Heligmosomoides polygyrus* (H.p.). *Lyz2^ΔOgt^* mice were already more susceptible to primary (1°) infection, harboring increased numbers of adult worms compared to *Ogt^flox^* mice, however resistance to 2° H.p. infection was not affected, indicating compensation by presence of a memory Th2 response (**Figure 2A**). At d7 of 1° infection, concordant with the emergence of larvae into the lumen and the onset of the initial Th2 response, *ΔOgt* macrophages within the small intestinal lamina propria (SILP) had impaired RELMα and PD-L2 expression compared to *flox* controls (**Figure 2B & S1A**). These data therefore confirm an *in vivo* role for O-GlcNAcylation in driving optimal AAM polarization and anti-helminth type 2 immunity.

**Figure 2.**
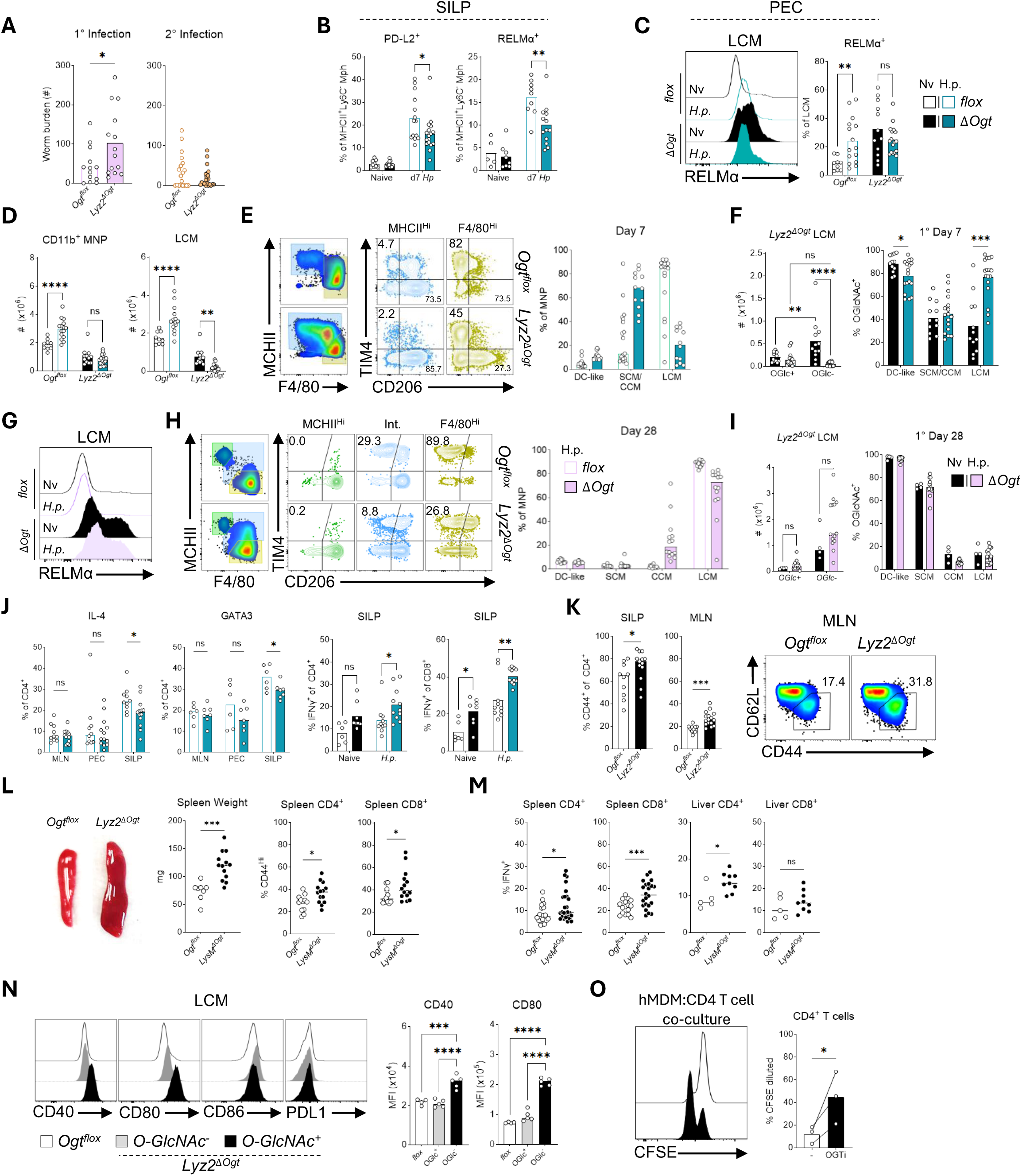
Macrophage O-GlcNAcylation is required for alternative activation in vivo and to control immune responses and during helminth infection. (**A**) Adult worm counts. Pooled from two experiments, n=6-8 mice/group (1°) or three experiments, n=5-9 mice/group(2°). (**B** & **C**) Alternative activation markers in mature macrophages at d7 H.p. infection from (B) duodenum/jejunum or (C) PEC. Pooled from three experiments, n=2-7 mice/group. (**D**-**F**) Analysis of peritoneal MNP. (D) Calculated MNP and LCM counts, (E) phenotypic analysis of SCM/LCM markers, and (F) quantification of O-GlcNAc^+^ MNP from *Lyz2^ΔOgt^* mice. Pooled from three experiments, n=4-7 mice/group. (**G**-**I**) Analysis of peritoneal MNP at d28 of infection. (G) Representative RELMα staining, (H) SCM/LCM marker expression in MNP and (I) quantification of O-GlcNAc^+^ in *Lyz2^ΔOgt^* mice. Naïve group from one experiment, infected pooled from two experiments, n=4-8 mice/group. (**J**) Cytokine expression by T cells following PMA/Ionomycin stimulation or native GATA3 at d7 of infection. GATA3 from one experiment, cytokines pooled from two experiments, n=4-6 mice/group. (**K**) Frequency of effector/memory CD4^+^ T cells in naïve mice. Pooled from two experiments, n=2-6 mice/group. (**L**) Representative spleen size, weight, and T cell activation. Pooled from three experiments, n=2-6 mice/group. (**M**) Intracellular IFNγ expression after PMA/Ionomycin. Pooled from six (spleen) and two (liver) experiments, n=2-6 mice/group. (**N**) Co-stimulatory marker expression of naïve LCM, representative of three experiments, n=4 mice/groups. (**O**) Proliferation of overnight pre-activated allogenic T cells after 5 days of co-culture at 1:2.5 ratio of hMDM:T cells, with hMDM differentiated normally or with daily treatment of 20µM OGTi. n=3 donors. Data presented as mean with biological replicates shown. Statistical comparison done with (A & M *spleen*) Welch’s or (J, K, L, M *liver*) standard unpaired t-test, (B-D, F & I) two-way ANOVA with Tukey’s correction or (F & I right, J *IFNγ*) Sidak correction for mean row comparisons, (N) one-way ANOVA with Tukey’s correction, or (O) ratio paired t-test.

H.p. infection also initiates an immune response in the peritoneal cavity. Here, *Ogt^flox^* macrophages increased RELMα expression at d7 of 1° infection in accordance with AAM polarization (**Figure 2C & S1B**). In contrast, *Lyz2^ΔOgt^* macrophages from naïve mice again exhibited elevated RELMα expression that was not further increased by infection. Similarly, total MNPs in the PEC failed to expand in *Lyz2^ΔOgt^* mice and showed a significant reduction in LCM (**Figure 2D**). The majority of MNPs in infected *Lyz2^ΔOgt^* mice instead resembled CCM, characterized by high MHCII and CD206, intermediate F4/80, and absence of the residence marker TIM4 (**Figure 2E**). Unlike IL-4c injection, macrophages within *Lyz2^ΔOgt^* mice showed that LCM were almost completely replaced by O-GlcNAc^+^ cells in response to H.p., suggesting additional signals from infection contribute to macrophage disappearance (**Figure 2F**). By d28 RELMα in *Ogt^flox^* mice returned to naïve levels, whereas *Lyz2^ΔOgt^* mice continued to show comparably high RELMα in both naïve and infected mice (**Figure 2G**). However, at this later timepoint, the MNP population resembled that of homeostatic mice for each respective genotype, where there remained a significant absence of TIM4^+^CD206^-^ mature LCM in the *Lyz2^ΔOgt^* strain (**Figure 2H**). Interestingly, O-GlcNAc levels also returned to that of naïve mice suggesting that, during resolution, repopulation of *Lyz2^ΔOgt^* LCM is driven by cells possessing residual O-GlcNAc or OGT activity that wanes over time (**Figure 2I**). Thus, O-GlcNAcylation is also needed for LCM maintenance during infection, but *Lyz2^ΔOgt^* LCM can recover during resolution, likely through monocyte recruitment.

T cell responses can be regulated by macrophages during infection^39,40^. We accordingly observed a decrease in intestinal IL-4 and GATA3 expression from CD4^+^ T cells at d7 of 1° infection, in *Lyz2^ΔOgt^* mice compared to *Ogt^flox^* mice, that was specific to the intestine (**Figure 2J & S1C**). This was not observed during challenge infection, supporting the presence of a memory Th2 response in both genotypes (not shown). Decreased Th2 cells paralleled patterns of increased IFNγ in both CD4^+^ and CD8^+^ T cells, indicating a localized shift from a Th2 to Th1 response. However, trends for increased IFNγ were already observed in uninfected *Lyz2^ΔOgt^* mice, along with increased CD4^+^CD44^Hi^ effector/memory cells in both the MLN and SILP, suggesting an altered homeostatic level of T cell activation (**Figure 2J & K**). *Lyz2^ΔOgt^* mice further presented with splenomegaly and systemically increased T cell activation and IFNγ expression (**Figure 2L & M**). We did not observe any differences in dendritic cell frequencies or activation in the spleen or SILP from either naïve or infected mice (**Figure S1D-G**). On the other hand, *<u>Ogt</u>*-deficient LCMs had higher co-stimulatory marker expression, suggesting better potential to directly support T cell activation (**Figure 2N**). Moreover, unstimulated human monocyte-derived M-CSF macrophages completely blocked proliferation of pre-activated T cells but were unable to do so when differentiated in the presence of OGTi (**Figure 2M**). Thus, macrophage O-GlcNAcylation is important for T cell homeostasis.

*Schistosoma mansoni* (S.m.) infection and B16.F10 melanoma were used as alternative models to assess to role of *Ogt* in AAM. Egg granuloma formation following S.m. infection was equivalent between both genotypes (**Figure S2A**). At 16 weeks post S.m. infection, all liver macrophages become monocyte-derived, even in wild-type mice, eliminating ontogeny as a confounder (**Figure S2B**). A significant decrease in AAM was still observed, despite high residual O-GlcNAc levels, which similarly correlated with reduced GATA3^+^ Th2 cells and increased IFNγ expression (**Figure S2C-E**). *Lyz2^ΔOgt^* also had significantly delayed tumor growth, which contained fewer pro-tumoral CD206^+^ macrophages, and increased IFNγ^+^ CD8 T cells (**Figure S2F-I**). Altogether, these data assert OGT is required for *in vivo* AAM polarization in multiple disease models and tissues.

### OGT promotes survival in response to environmental or immunological stimuli

As RELMα expression is a feature of monocyte-derived macrophages in the peritoneal cavity, the increase in RELMα^+^ LCM in unchallenged *Lyz2^ΔOgt^* mice indicated a baseline disruption in macrophage homeostasis. Concordantly, we found *Lyz2^ΔOgt^* mice had proportionally fewer LCM and more CCM (**Figure 3A & B**). Overall MNP numbers from the cavity were decreased in *Lyz2^ΔOgt^* mice, due to selective loss of LCM (**Figure 3B**). Only LCM fully lost O-GlcNAcylation in *Lyz2^ΔOgt^* mice, while a progressive decrease in GlcNAcylation from DC-like to SCM and CCM MNPs could be observed (**Figure 3C**). Combined with reduced TIM4 expression in LCM (**Figure 3D**), these data point towards loss of LCM maintenance that is compensated by repopulation by monocyte-derived cells in *Lyz2^ΔOgt^* mice.

**Figure 3.**
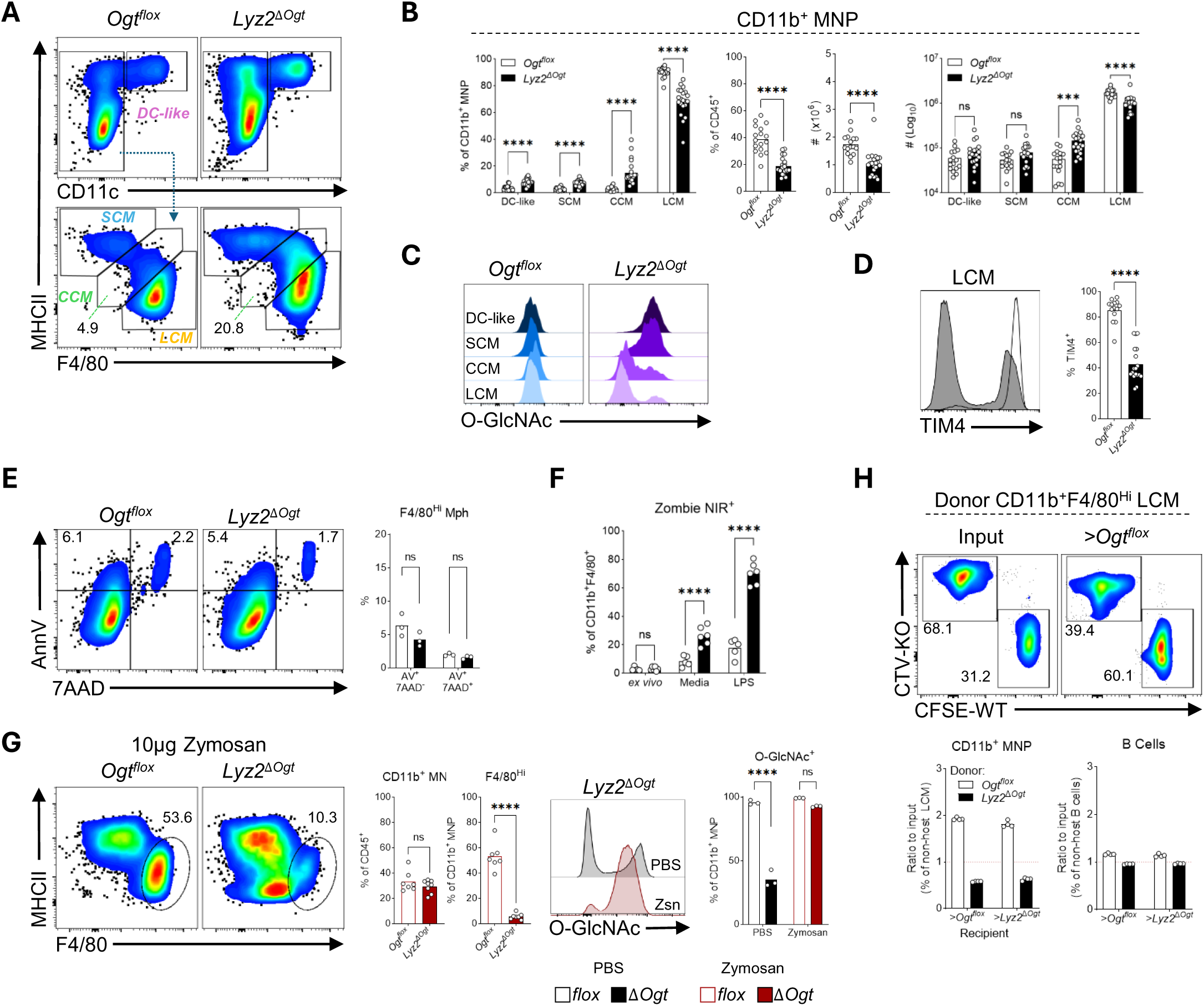
*Ogt*-deficient macrophages are susceptible to stress-induced cell death. (**A** & **B**) Characterization of peritoneal MNP composition. (A) Gating and (B) quantification of MNP populations. Six experiments pooled, n=2-4 mice/group. (**C**) Representative O-GlcNAc staining for MNP populations as gated in (B). (**D**) Expression of residency marker TIM4. Pool of four experiments, n=2-4 mice/group. (**E**) Cell viability of LCM directly *ex vivo* determined by Annexin V and DNA stain 7AAD. Representative of two experiments, n=3 mice/group. (**F**) Viability of *ex vivo* cultured LCM unstimulated or treated with 100ng/ml LPS for 5 hours. Representative of two experiments, n = 5-6 mice/group. (**G**) Analysis of MNP population d3 after i.p. zymosan injection. Representative or pooled from two experiments, n=3-4 mice/group. (**H**) Recovery of labelled LCM 24 hours after total PEC was labelled and transferred intraperitoneally to recipients. 1 experiment performed, n = 4 (*Ogt^flox^*) or 7 (*Lyz2^ΔOgt^*) donors pooled and 4 recipients/group. Data presented as mean with biological replicates shown. Statistical comparison done with (A, D, E, H) unpaired t-test, (A) RM two-way ANOVA row comparison with Sidak correction, or (F, H) two-way ANOVA with Tukey’s correction.

We first hypothesized that *Ogt* is required for survival of LCM. However, we did not observe any differences in viability directly *ex vivo*, based on DNA and Annexin V staining (**Figure 3E**), suggesting *Lyz2^ΔOgt^* LCM survive similarly, at least in the absence of perturbation. Previous work has shown *Ogt* prevents cell death in response to bacterial stimulation^21^. Similarly, we found that LPS caused significant cell death in *Ogt-*deficient compared to sufficient LCM. We however observed notable cell death from culture alone in *Lyz2^ΔOgt^* LCM, whereas viability staining was again comparable prior to culture (**Figure 3F**). To assess cell death *in vivo*, we injected a low dose of zymosan, which triggers incomplete disappearance of resident LCM in C57BL/6 mice^29^. In *Lyz2^ΔOgt^* mice, we instead observed a complete loss of F4/80^Hi^ LCMs, and those remaining were all TIM4^-^ (**Figure 3G**). Accordingly, less than 10% of total *Lyz2^ΔOgt^* MNPs remained O-GlcNAc^+^, confirming lost LCM are replenished by temporarily OGT-sufficient cells.

To directly test the role of OGT in maintaining macrophage residency we co-transferred dye-labelled PEC from *Ogt^flox^* or *Lyz2^ΔOgt^* mice into *Ogt^flox^* recipients. Within 24 hours of transfer, we observed a rapid loss of *Lyz2^ΔOgt^* LCM, indicated by the shifted ratio of recovered *flox* to *ΔOgt* LCM compared to the input ratio (**Figure 3H**). This was specific to LCM as the ratio of B cells did not change. A similar transfer into *Lyz2^ΔOgt^* recipients revealed a comparably shifted ratio, suggesting the loss of *Ogt*-deficient LCM is driven primarily by a cell intrinsic mechanism. Together, these findings suggest the loss of LCM in *Lyz2^ΔOgt^* mice can be explained, in part, by sensitization to cell death which is accentuated during immunological stress.

### O-GlcNAcylation enforces macrophage quiescence while preventing a senescence

To explore potential additional mechanisms leading to loss of LCM maintenance, we performed single-cell RNA sequencing on the total PEC from *Ogt^flox^* and *Lyz2^ΔOgt^* mice. Analysis and annotation of MNPs revealed a cluster of altered LCM from *Lyz2^ΔOgt^* mice transcriptionally distinct from normal LCM (**Figure 4A & Figure S3**). Except for cDC1, all MNP formed a continuum in UMAP space, indicative of a cell development trajectory, supported by RNA velocity analysis which recapitulated previously defined serous cavity macrophage subset relationships^36^. The flux of CCM towards tissue residency appeared to hold true for both genotypes, however within the altered LCM, velocity vectors indicated a flux into the proliferation (G2M Phase) cluster (**Figure 4B**). Scoring MNP against foundational LCM and SCM ImmGen datasets revealed that altered LCM scored between CCM and LCM, indicating the altered LCM share many features of bona-fide LCM but few can completely adopt the full LCM program (**Figure 4C**). Nonetheless, GATA6 protein expression was unchanged and LCM from *Lyz2^ΔOgt^* mice and correspondingly, still possessed high predicted GATA6 regulon activity (**Figure 4D**).

**Figure 4.**
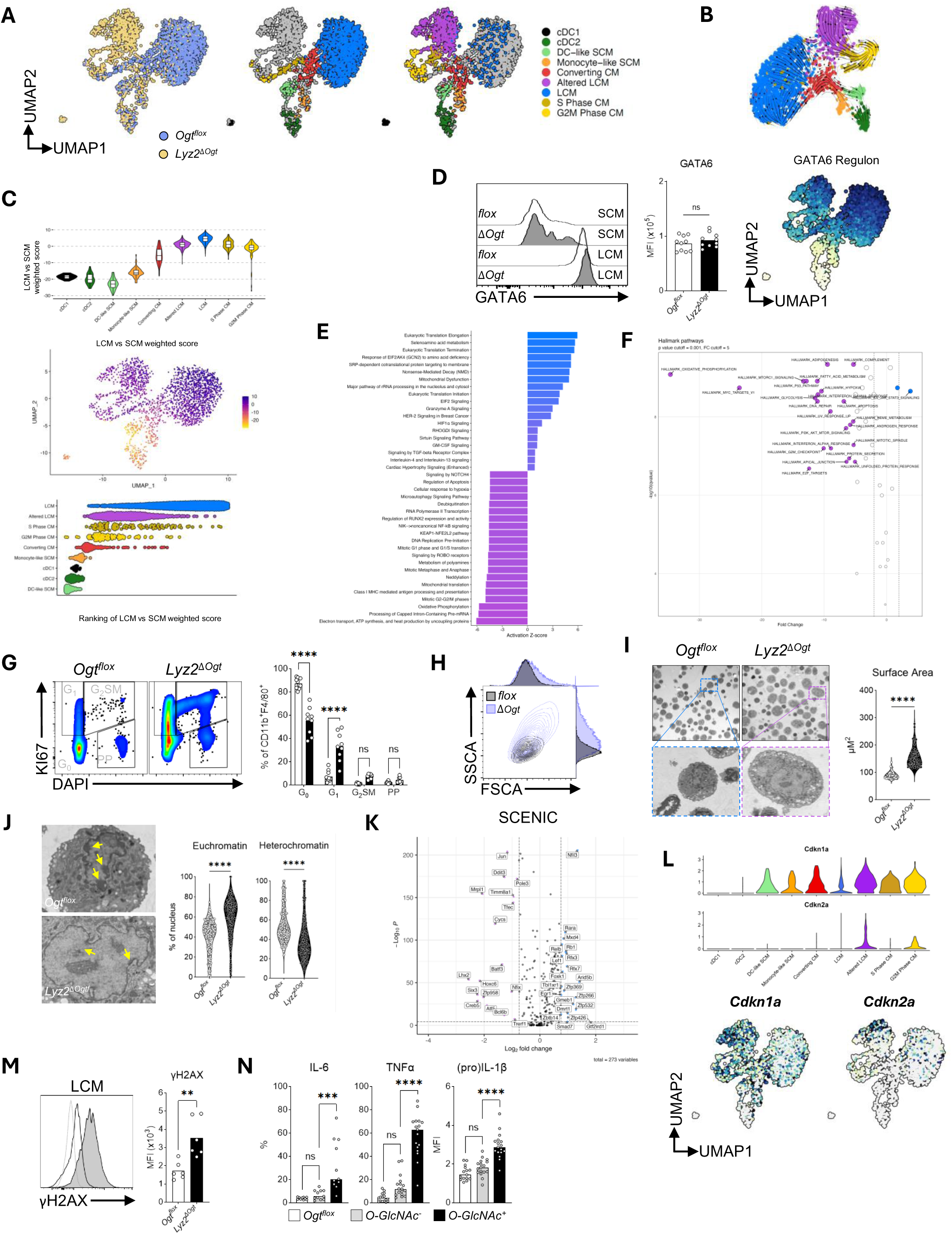
*Ogt* is required to prevent a senescent-like phenotype in macrophages. (**A**) UMAP and MNP clusters generated from differential gene expression of single-cell RNA sequencing data subset from an initial 12,045 singlets from total PEC after QC and filtering, n=3 mice/group. (**B**) RNA velocity analysis. (**C**) Calculation, projection and ranking of SCM-LCM scores, constructed from Immunological Genome Project datasets. (**D**) GATA6 protein expression in LCM and GATA6 transcriptional regulon score. GATA6 expression pooled from two experiments with n = 3-7mice/group. Unpaired two-tailed t-test performed, ns = not significant. (**E** & **F**) Top differentially expressed pathways using (E) IPA or (F) Hallmark pathways, between normal and altered LCM. (**G**) Cell-cycle analysis of LCM. Data pooled from three experiments, n=2-4 mouse/group. (**H**) Representative forward and side scatter plot of peritoneal LCM. (**I**) Electron microscopy scan and quantification of cell size by AI identification and calculation. Top 20% of largest identified macrophages used for calculation, n = 2 mice, filtered for largest 88 cells/genotype. (**J**) EM images of peritoneal macrophage nuclei. Surface area of heterochromatin and euchromatin determined by machine learning identification and calculation. *Ogt^flox^* = 300, *Lyz2^ΔOgt^* = 220 cells. (**K**) Predicted differential SCENIC transcription factor regulons. (**L**) Differential expression of cyclin dependent kinases between MNP populations. (**M**) DNA damage staining in naïve LCM. Pooled from two experiments, n=3 mice/group. (**N**) Cytokine expression in unstimulated LCM cultured *ex vivo* with brefeldin A and monensin for five hours. Three or four experiments pooled, n=3-6 mice/group. Unless otherwise indicated datapoints are biological replicates. Statistical comparison done with (C, I, J & M) unpaired t-test, (G) RM two-way ANOVA row comparison with Sidak correction, or (N) one-way ANOVA with Tukey’s correction.

In light of similar steady-state viability and similar, though not identical, cellular identity of LCM from *Ogt^flox^* and *Lyz2^ΔOgt^* mice, we asked whether O-GlcNAcylation controls proliferative capacity and renewal. Pathway analysis accordingly indicated altered LCM had a highly dysregulated cell cycle, accompanied by changes in translational and transcriptional regulation, as well as metabolism (**Figure 4E & F**). In line with CCM flux towards the proliferative clusters, and significantly more *Lyz2^ΔOgt^* MNP transcriptionally falling in the proliferative cluster itself, *Lyz2^ΔOgt^* LCM were found predominantly outside the G0 quiescent phase when stained for DNA and Ki67 (**Figure 4G & S3L**). Yet, *Ogt*-deficient LCM were larger and more granular than normal LCM (**Figure 4H**), and electron microscopy confirmed an approximate doubling of cell size (**Figure 4I**). Furthermore, MNP from *Lyz2^ΔOgt^* mice substantially lacked visible heterochromatin (**Figure 4J**), consistent with heightened transcriptional activity and cell growth. Combined with the homeostatic reduction of *Lyz2^ΔOgt^* LCM and their inability to expand directly in response to IL-4 (**Figure 1I-K**), these data support *Ogt*-deficiency increases cell cycle progression, and concordant cellular growth, without the ability to complete it.

The dysregulated cell-cycle and loss of LCM observed in *Lyz2^ΔOgt^* mice is reminiscent of *Bhlhe40* deletion, which also showed a defective proliferative response to IL-4^41^. Correspondingly, the transcriptional signature of altered LCM from *Lyz2^ΔOgt^* mice was highly similar to *Bhlhe40-*deficient LCM (**Figure S4A**), indicating that Bhlhe40 might be downregulated or targeted by O-GlcNAcylation. However, we found no loss of BHLEH40 protein in LCM from *Lyz2^ΔOgt^* mice (**Figure S4B**), or any evidence that BHLEH40 was directly O-GlcNAcylated (**Figure S4C & D**), possibly pointing towards indirect regulation of BHLEH40 activity by O-GlcNAcylation.

Since increased cell size without dividing is a hallmark of cellular senescence, we questioned whether the replicative dysfunction induced by *Ogt*-loss is associated with senescence in LCM. Supportively, despite *Lyz2^ΔOgt^* LCM engaging the cell cycle, multiple pathways and transcriptional regulons enriched in altered LCM aligned with a senescent-like phenotype. For instance, Hallmark pathways dominant in altered LCM included: IFNα response, DNA repair, UV response, P53 pathway, unfolded protein response and protein secretion (**Figure 4F**). Using SCENIC regulon analysis (**Figure 4K**), altered LCM also had higher predicted activation of *Jun*, recently identified as part of a core senescence signature^42^, *Irf7*, a master regulator of type 1 IFN responses that has been linked to the DNA damage response^43,44^, and *Ddit3* (CHOP), central to cell stress induced by the unfolded protein response^45^. Indeed, the senescence-defining genes p21-cip1(*Cdkn1a*) and p16-ink4a (*Cdkn2a*) were more highly expressed in altered LCM (**Figure 4L**). We confirmed that *Lyz2^ΔOgt^* LCM further exhibited increased DNA damage (**Figure 4M**), as well as spontaneous secretion of inflammatory cytokines commonly associated with the senescence associated secretory phenotype (**Figure 4N**). Therefore, the loss of *Ogt* in macrophages strongly aligns with the acquisition of a senescent state^46^.

Altogether, we find O-GlcNAcylation is required to maintain the homeostatic peritoneal macrophage population through a combination driving self-renewal, preventing immune-senescence and promoting survival in response to diverse stimuli.

### Dysregulated metabolism in **Δ**Ogt LCM leads to glucose dependent ROS production

Senescent cells additionally frequently present with metabolic dysfunction, and our scRNA-seq dataset suggested highly divergent metabolic programming in altered LCMs (**Figure 4E & F**). In confirmation, LCMs from *Lyz2^ΔOgt^* mice had significantly higher Myc expression, and phosphorylated S6, pointing towards elevated mTOR signaling (**Figure 5A**). *Lyz2^ΔOgt^* LCM also displayed higher pACC, representing active AMPK signalling, however there was a greater relative increase in pS6, suggestive of a shift towards anabolic metabolism (**Figure 5A**).

**Figure 5.**
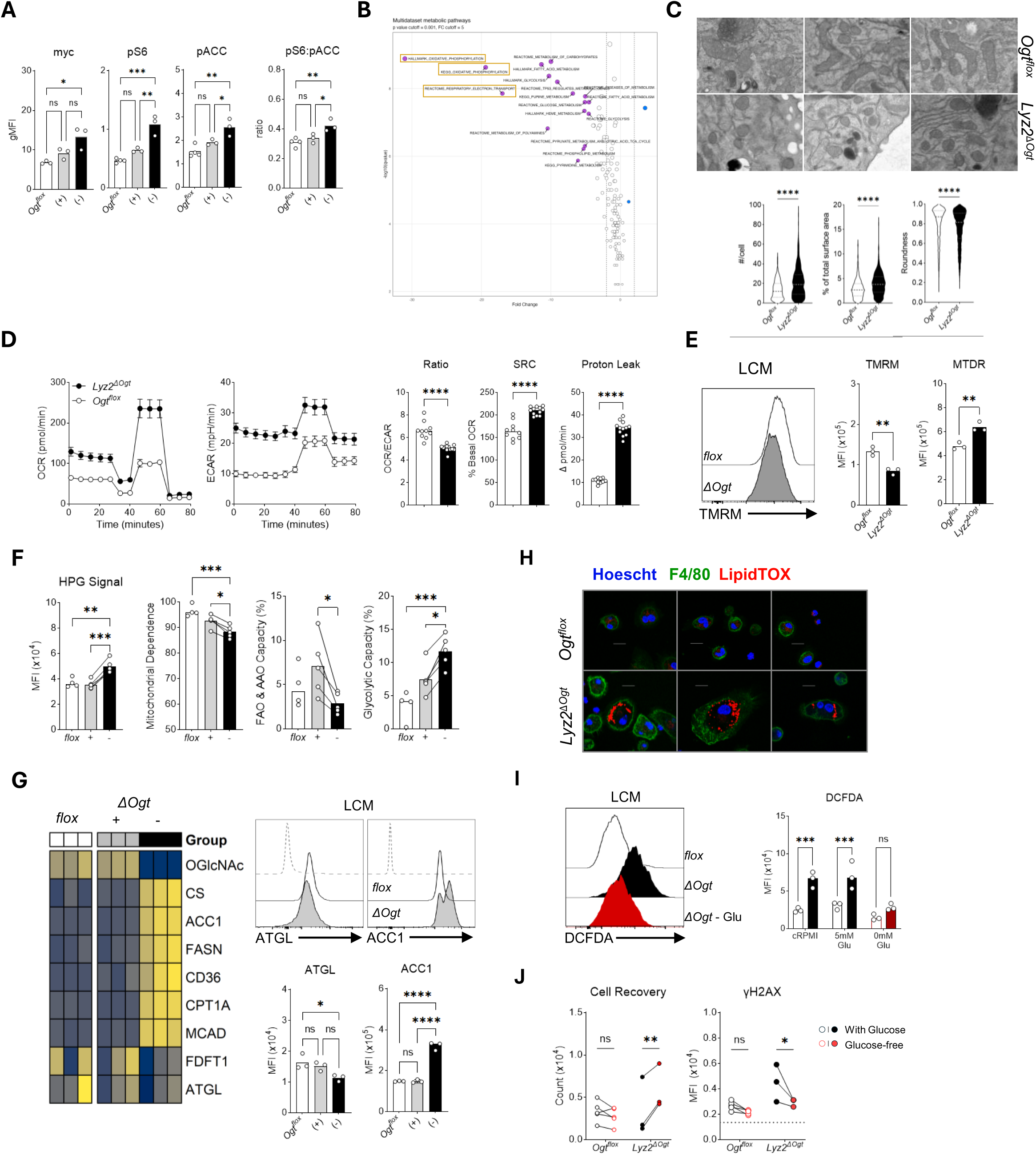
Mitochondrial metabolism, lipid storage and ROS production are regulated by O-GlcNAcylation in macrophages. (**A**) Expression of total Myc, and mTOR and AMPK signaling by proxy of phosphorylated targets S6 and ACC1. Representative of two experiments, n=3 mice/group. (**B**) Concatenated analysis of metabolic pathways from HALLMARK, KEGG and REACTOME lists using differential expression from scRNA-seq between normal and altered LCM. (**C**) Example images of mitochondria and machine learning quantification of relative surface area, roundness and counts per cell, *Ogt^flox^* = 5284 and *Lyz2^ΔOgt^* = 6951 mitochondria analyzed. (**D**) Seahorse mitochondrial stress test performed on sorted CD11b^+^F4/80^Hi^MHCII^-^ LCM pooled from 2-3 mice/group, and plated in technical replicates. (**E**) TMRM and MitoTracker DeepRed detection, representative of three experiments, n=3 mice. (**F**) L-Homopropargylglycine (HPG) incorporation as a proxy for translation, and metabolic dependencies assessed following treatment with 2-DG or oligomycin. Represents one experiment, n=4-5 mice/group. (**G**) Metabolic flow cytometry expression of enzymes involved in lipid metabolism. One experiment performed, n-3 mice/group. (**H**) Confocal images of neutral lipid staining in peritoneal macrophages. Images are from separate mice from one of two experiments. (**I**) Total ROS detection in LCM from total PEC incubated with DCFDA *ex vivo*, representative of 2 experiments, n=3 mice/group. (**J**) Total CD11b^+^F4/80^Hi^ events after five hours of *ex vivo* culture and MFI of yH2AX within live cells. One experiments with 3-5mice/group. Data presented as mean with biological replicates except for C & D. Statistical comparison done using (A, F and G) one-way ANOVA with Tukey’s correction, (C-E) unpaired t-test, or (I & J) two-way ANOVA row comparison with Sidak correction.

We found that oxidative phosphorylation (OxPhos) was consistently the top enriched pathway in the altered compared to normal LCMs (**Figure 4D, E & 5B**). Electron microscopy image quantification validated that *Lyz2^ΔOgt^* macrophages had higher mitochondrial content (**Figure 5C**). Furthermore, mitochondria from *Lyz2^ΔOgt^* LCM had a more elongated morphology characteristic of fusion, as well as more disorganized cristae (**Figure 5C**). These changes in mitochondria were associated with an overall increase in oxygen consumption in sorted *Lyz2^ΔOgt^* compared to *Ogt^flox^* LCM (**Figure 5D**). *Ogt-*deficient LCM also exhibited elevated aerobic glycolysis, such that the ratio of oxygen consumption compared to extracellular acidification was reduced compared to normal LCM, pointing towards a preferential shift towards glucose dependence (**Figure 5D**). LCM sorted from *Lyz2^ΔOgt^* mice also possessed higher spare respiratory capacity (SRC), as well as proton leak. These observations corresponded to a reduction in overall membrane potential, determined by TMRM staining, despite MitoTracker staining again indicating *Lyz2^ΔOgt^* LCM have more mitochondria (**Figure 5E**), concordant with a high rate of ATP generation to stabilize the protonmotive force. Further analysis of metabolic dependencies for translation using click-chemistry based detection of L-Homopropargylglycine (HPG) incorporation^32,47^, paired with SCENITH^48^, showed higher translational activity in altered LCM and decreased mitochondrial dependence (**Figure 5F**).

As we observed a decreased fatty acid dependence in altered LCM (**Figure 5F**), we assessed enzyme expression for lipid pathways by flow cytometry. Only the lipase ATGL and cholesterol biosynthesis enzyme FDFT1 were not increased in O-GlcNAc^-^ LCM (**Figure 5G**). We also found *Ogt*-deficiency led to decreased expression of the lipase *Lpl*, but increased expression of several fatty acid binding proteins (FABPs) involved in intracellular lipid trafficking and storage, and the lipid droplet stabilizer perilipin 1 (*Plin1*) (**Figure S5A**). Accordingly, *Lyz2^ΔOgt^* LCM, as well as BMDM, accumulated neutral lipids, forming large droplets within a short time of *ex vivo* culture (**Figure 5H & S5B**), corroborating altered LCM divert lipids towards storage. Lipid droplet formation can be a strong indication of reactive oxygen species (ROS) production. We therefore hypothesized that the high mitochondrial activity and associated proton leak of *Ogt*-deficient LCM contributed to increased ROS as a driver for DNA damage and senescence. DCFDA staining supportively showed that LCM from *Lyz2^ΔOgt^* mice produced significantly more ROS (**Figure 5I**), which was recapitulated by the addition of OGTi to wildtype BMDM (**Figure S5C**). Elevated ROS production was almost entirely dependent on glucose (**Figure 5I**), and glucose withdrawal over short term culture reduced the overall loss of peritoneal macrophages, concomitant with a reduction in DNA damage (**Figure 5J**).

These observations underscore O-GlcNAcylation as a central regulator of macrophage metabolism. As such, OGT deletion leads to hyperactive, glucose-dependent, mitochondrial respiration to support increased energetic demands from unrestrained growth. Subsequently ROS production may then be a key driver of the senescent-like state by damaging exposed DNA.

### Dynamic regulation of O-GlcNAcylation during LCM differentiation

At steady-state, the majority of tissue-resident macrophages remain quiescent, whereas monocyte-derived populations are more readily found in an active cell cycle, despite the non-cycling nature of blood monocytes^27,29,49,50^. Thus monocyte-macrophages must actively engage the cell cycle during tissue entry and exit again following differentiation. Considering the unrestrained metabolic activity in LCM due to loss of *Ogt*, we predicted dynamic O-GlcNAcylation would be required to dampen proliferative and metabolic activity during residence formation. Accordingly, we found O-GlcNAcylation from *Ogt^flox^* mice was elevated in CCM and LCM compared to SCM (**Figure 6A**). Indeed, expression of metabolic proteins between *Ogt^flox^* and *Lyz2^ΔOgt^* mice was similar for SCM, but significantly higher in O-GlcNAc^-^ LCM compared to O-GlcNAc^+^ LCM (**Figure 6B**). As previously reported^32^, CPT1A and GLUT1 expression was reduced in normal LCM compared to SCM, which paralleled a transient increase in mTOR and shift towards AMPK signaling (**Figure 6B**). In *Lyz2^ΔOgt^* mice, metabolic target expression instead increased in LCM relative to SCM, paralleling continually elevated mTOR activation (**Figure 6C**). In concordance with this metabolic transition, a large proportion of *Ogt^flox^* SCM and CCM were found actively in the cell cycle relative to LCM, whereas *Lyz2^ΔOgt^* LCM possessed Ki67 accessibility similar to CCM (**Figure 6D**). Principal component (PC) analysis of MNP from our scRNA-seq dataset correspondingly revealed similarity in PC1 between altered LCM and proliferative cells, suggesting a residual cell cycle signature in altered LCM (**Figure 6E**).

**Figure 6.**
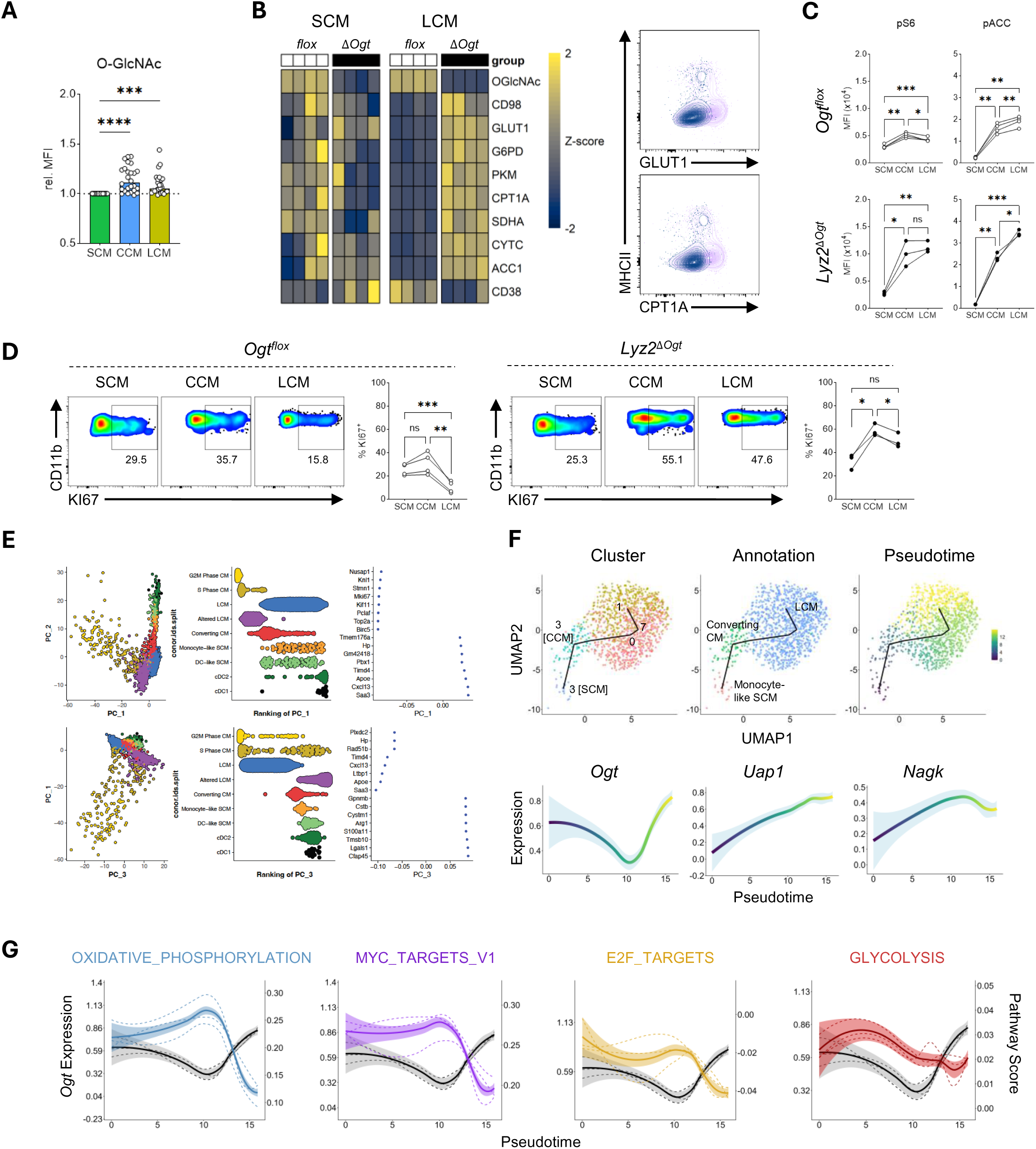
Restoring metabolic and cell-cycle quiescence requires O-GlcNAcylation during macrophage differentiation. (**A**) Detection of O-GlcNAc normalized to SCM, retrospectively pooled from eight experiments, n = 2-4 mice/group. (**B**) Row-scaled heatmap of metabolic protein expression detected by flow cytometry, and representative staining for GLUT1 and CPT1A in total CD11b^+^CD11c^-^ MNP, n = 4 mice/group, representative of two experiments. (**C**) mTOR and AMPK signaling by proxy of phosphorylated targets S6 and ACC1. Representative of two experiments, n=3-4 mice/group. (**D**) Frequency of MNP populations outside G0 of the cell-cycle. Representative of three experiments, n=3-4 mice/group. (**E**) Projection of MNP clusters from scRNA-sequencing onto principle component analysis plots, corresponding ranking of clusters and top defining genes for PC1 and PC3. (**F**) TSCAN scRNA-seq pseudo-time trajectory of *Ogt^flox^* MNP and corresponding expression of HBP genes according to trajectory. (**G**) Calculated scores of HALLMARK pathway gene expression compared to *Ogt* along calculated pseudo-time in (F). Statistics for (A, C & D) calculated using one-way RM ANOVA and Tukey post-hoc test.

To provide additional evidence of temporal cell-cycle and metabolic regulation by O-GlcNAcylation during residency formation, we performed transcriptional pseudotime analysis. In *Ogt^flox^* MNP, both *Ogt* and *Uap*, a key enzyme in the HBP, exhibited temporal regulation with increasing expression towards mature LCM, whereas the UDP-GlcNAc salvage gene *Nagk* showed a late decrease (**Figure 6F**). We calculated Hallmark pathway scores over the course of the trajectory and found a clear inverse correlation between *Ogt* expression and OxPhos, Myc and E2F targets, and to a lesser degree glycolysis, such that pathway scores were lowest when *Ogt* expression was highest (**Figure 6G**). Cell-cycle and metabolism are closely connected, and thus difficult to segregate, however, non-proliferative BMDM still acquired several overlapping phenotypes when comparing *Lyz2^ΔOgt^* versus *Ogt^flox^* cells, without apparent changes in cell-cycle (**Figure S5E-F**). Therefore O-GlcNAcylation may control metabolism upstream, or independent of proliferation. These observations together support a model in which LCM differentiation is accompanied by dynamic metabolic programming and cell cycle kinetics, ending with a re-acquisition of quiescence that is unattainable in the absence of O-GlcNAcylation.

### O-GlcNAcylation is required for macrophage homeostasis across organs

Lastly, we questioned the importance of O-GlcNAcylation for other tissue resident macrophage populations. Surprisingly, compared to peritoneal cavity, macrophages from the lung, liver and intestines showed high remaining levels of O-GlcNAc, with alveolar macrophages (AlvM) showing little difference between those from *Lyz2^ΔOgt^* or *Ogt^flox^* mice (**Figure 7A**). The relative loss in O-GlcNAcylation appeared inversely related to O-GlcNAc levels in naïve *Ogt^flox^* mice. Despite the remaining levels of O-GlcNAc, *Lyz2^ΔOgt^* mice had significantly fewer macrophages in the lung, which was specifically due to a loss of SiglecF^+^ AlvM (**Figure 7B**). Kupffer cells (KC) from the liver were also numerically reduced, and fewer expressed TIM4 (**Figure 7C**). Despite similar loss of resident macrophages in these tissues, O*gt-*deficiency differently affected their size and granularity. AlvM from *Lyz2^ΔOgt^* mice had increased granularity but comparable size, whereas for KC both increased in parallel (**Figure 7D**). Following IL-4c injection, both *Ogt*-sufficient AlvM and KC expanded in numbers compared to PBS, whereas *Ogt*-deficient cells failed to do so (**Figure S6A**). In this context AlvM rapidly lost O-GlcNAcylation, suggesting steady-state O-GlcNAc levels are largely due to the inactivity of OGA. Accordingly, *Oga* expression was also reduced in peritoneal *Lyz2^ΔOgt^* LCM to similar levels as *Ogt* (**Figure S6B**). The reduction in homeostatic AlvM may then be a consequence of blocked proliferation, rather than cell death. KC on the other hand showed a modest decrease in O-GlcNAc^+^ cells, which were increased following IL-4 exposure, suggesting a contribution of cell death and replacement (**Figure S6B**).

**Figure 7.**
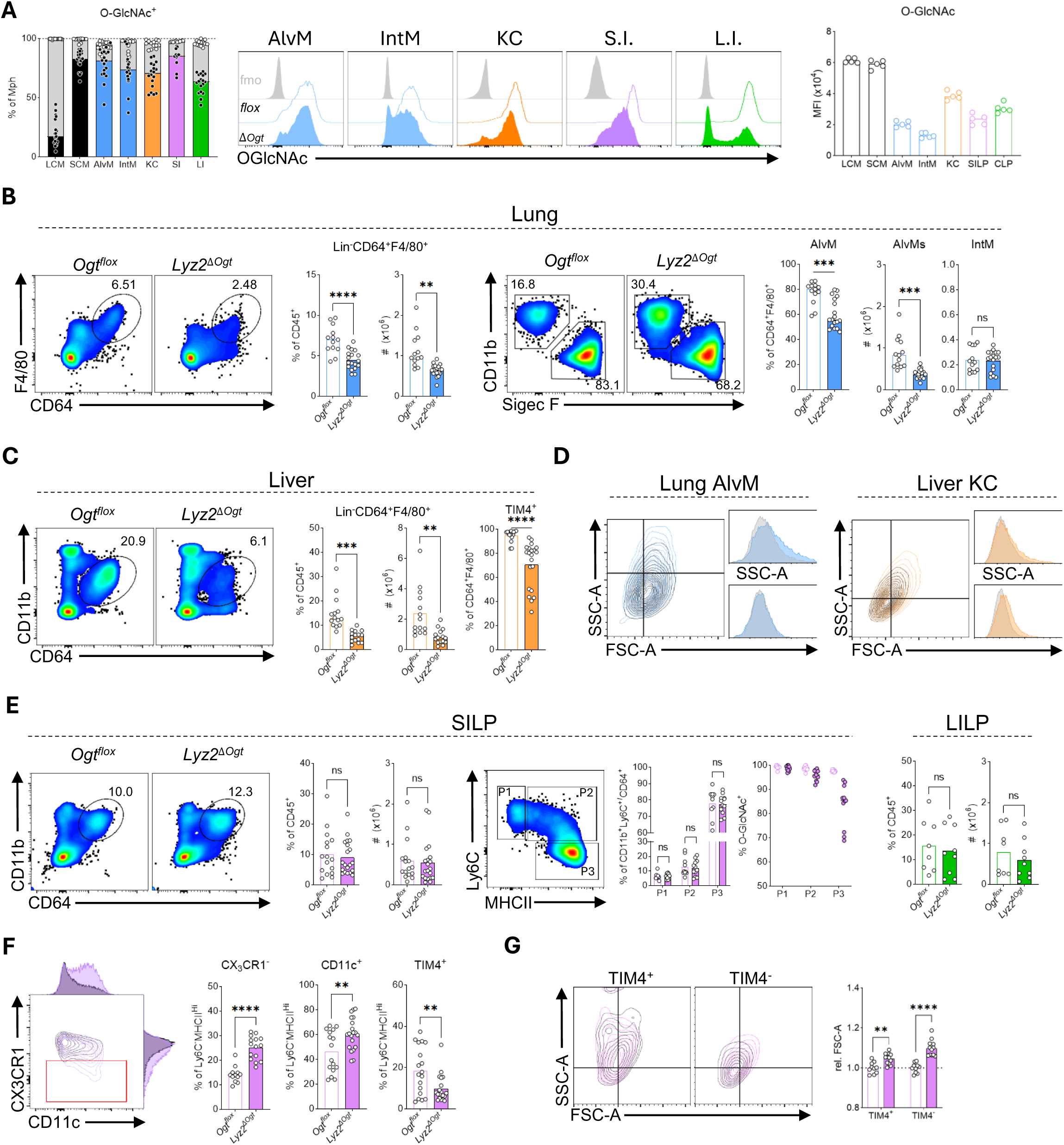
Macrophage requirements for O-GlcNAcylation vary according to tissue site. (**A**) Detection of residual O-GlcNAc across tissue macrophage populations and relative O-GlcNAc levels in *Ogt^flox^* mice. SILP pooled from three, otherwise five experiments, n=2-6 mice/group. O-GlcNAc MFI in *Ogt^flox^* represents one of three experiments, n=5. (**B** & **C**) Characterization of (B) lung and (C) liver macrophages. Representative detection of total macrophages from live, single, CD45^+^ cells and quantification of Lin^-^CD64^+^F4/80^+^ cells or respective tissue subsets. Pooled mice from five experiments for each tissue, n = 2-6 per group. (**D**) Representative scatter plots and histograms of lung AlvM or liver KC. (**E**) Characterization of small and large intestine macrophages. Representative detection of total macrophages from live, single, CD45^+^ cells, quantification of Lin^-^CD64^+^/Ly6C^+^ cells, frequency of monocyte waterfall populations and their corresponding O-GlcNAc levels. Pooled mice from five (SILP) or three (CLP) experiments, n = 2-5 per group. (**F**) Example staining and analysis of CD11c and CX_3_CR1, or TIM4 expression in intestinal macrophages, pooled from five experiments. (**G**) Representative size scatter plots and quantification of TIM4^+^ and TIM^-^ populations, pooled from three experiments. Statistics calculated using unpaired two-tailed t-test with Welch’s correction applied in cases of unequal variance.

The same results were observed using *Itgax*-cre *Ogt^flox^* mice, to target CD11c-expressing cells, particularly for AlvM, both regarding O-GlcNAc levels and cell numbers (**Figure S6C**). Peritoneal LCM were the exception, which do not express CD11c if embryonically seeded. However, there was a reduction in DC-like MNP and SCM consistent with their current or historical expression of CD11c^38^. As *Ogt* is found on the X chromosome, and macrophage replenishment differs between male and female mice^38^, we retrospectively analyzed experiments according to sex (**Figure S7**). Differences tended to be stronger in males, consistent with their known faster replenishment rates, but overall data were comparable using either sex of *Lyz2^ΔOgt^* mice. Similarly, peritoneal LCM decreased with aging at similar rates in both genotypes (**Figure S7A**).

In contrast to most tissues, the intestines contain a large population of continually replaced monocyte-derived macrophages^49^. Small intestine and colonic macrophage numbers were comparable between *Ogt^flox^* and *Lyz2^ΔOgt^* mice (**Figure 7E**). While there was no observable change in the monocyte waterfall between genotypes, O-GlcNAc levels were reduced progressively from monocytes (P1) to intermediate (P2) and mature MHCII^+^ macrophages (P3), similar to peritoneal macrophages, likely due to delay between *Ogt* loss and complete O-GlcNAc removal by OGA. Overall numbers were unchanged, however fewer MHCII^+^ macrophages from the SILP expressed CX_3_CR1, a marker of maturation in the intestine, while more expressed the inflammatory marker CD11c (**Figure 7F**). Similarly, TIM4^+^ embryonically derived macrophages were reduced, but the difference was interestingly more specific to male mice (**Figure 7F & S7B**). Furthermore, *Ogt*-deficiency had a greater effect on TIM4^-^ macrophages, which possessed a more significant size increase than TIM4^+^(**Figure 7G**), suggesting monocyte-derived macrophages in the intestine have more reliance on O-GlcNAcylation.

## DISCUSSION

We characterized tissue macrophage populations of mice with restricted O-GlcNAcylation via deletion of the modifying enzyme OGT. We report a marked deficiency in IL-4 responsiveness and in residency of macrophages across organs, linked to acquisition of senescent-like state, characterized by hyper-elevated ROS production, exposed DNA and subsequent DNA damage. Our findings present O-GlcNAcylation as novel and central regulatory node for establishing residency through metabolic adaptation and promoting cellular quiescence, primarily following a monocyte differentiation program.

Our data support an inability to self-renew as a primary cause for reduced macrophage numbers in *Lyz2^ΔOgt^* animals. O-GlcNAcylation is a well-established rheostat for positively regulating cell cycle entry and progression^35^. Correspondingly, we find macrophages deficient for *Ogt* have uncontrolled cell cycle entry, yet display several signs of failed division, namely dramatic enlargement relative to normal LCM and inability to expand in response to IL-4. Presumably, the observed increase in steady-state replacement of *Lyz2^ΔOgt^* LCM by monocyte-derived cells serves as a compensatory response for the lack of renewal. We do not find signs of increased cell death rates at steady state, although *Ogt*-deficient LCM more rapidly die following perturbation *in vivo* or *ex vivo*, which in the former case results in accelerated reconstitution of the niche by monocyte derived cells. This largely aligns with a report showing, *Lyz2^ΔOgt^* BMDM are more susceptible to necroptosis, but only in the context of LPS stimulation^21^. Thus, is it also possible that random daily or circadian events that may trigger stress or inflammation, such as fighting or feeding, contributes to macrophage loss and replacement.

In addition to cell renewal, our data suggest O-GlcNAcylation is also critical to establish macrophage metabolic and immunological quiescence following maturation from blood monocytes. Monocytes are typically non-proliferative, however enter the cell cycle during macrophage differentiation following tissue entry, and ultimately shift again toward a low number of quiescent cycling resident cells^27,29,49,50^. Accordingly, dampened mTOR activity is required for residence formation, partially by reducing hyperinflammatory stimulation and cell death^31,51^. We found cellular metabolism closely follows the same trajectory towards cell cycle quiescence through stages of monocyte to macrophage differentiation. Importantly, this metabolic transition that was highly dependent on O-GlcNAcylation as macrophages from *Lyz2^ΔOgt^* mice displayed a hypermetabolic state accompanied by elevated mTOR signaling. Furthermore, constitutive mTOR signaling is a driver of cellular senescence^52^. In line with recent definitions^46^, we find *Lyz2^ΔOgt^* macrophages acquire a senescent-like phenotype, including lipid accumulation aberrant ROS production, and DNA damage. Senescence in these macrophages may be initially triggered, in part, by an inability to down-regulate mTOR signaling during differentiation. Although similar observations have been made in other cell types^53,54^, this has not been reported in macrophages, likely due to their non-proliferative nature *in vitro*. Interestingly, recent works have linked IL-4 signaling to LCM formation^36^ and the prevention of senescence by promoting DNA repair^55^. Hence, O-GlcNAcylation of STAT6 may have dual responsibilities in both promoting AAM polarization as well as mitigating DNA damage induced senescence. Interestingly, BHLHE40 was found to be a direct transcriptional repressor of cell cycle proteins^41^, and we found *Bhlhe40* deficient LCM to be highly similar to *Ogt* deficiency but no evidence that BHLHE40 was targeted for O-GlcNAcylation. Whether BHLHE40 is an indirectly regulated O-GlcNAcylation of upstream targets or binding partners requires further study.

Accordingly, we find a critical role for macrophage O-GlcNAcylation in IL-4 responsiveness. Several pieces of evidence point towards O-GlcNAcylation as required for AAM differentiation: the HBP is upregulated in response to IL-4^16,19,20^, STAT6 contains multiple O-GlcNAc sites^17,18^, and OGT inhibitors block STAT6 transcriptional activity^17,18^. In epithelial cells O-GlcNAcylation controls STAT6 nuclear interactions^18^, rather than trans-activation, consistent with our data showing normal STAT6 tyrosine phosphorylation. However, we predict STAT6 independent mechanisms also contribute. Cellular metabolism is also central to macrophage polarization^8,10^, thus the resulting changes in nutrient sensing, or usage, by disrupted O-GlcNAcylation are likely impactful. Conversely, glucose, fatty acids and glutamine all form a nexus at the HBP and may promote AAM through O-GlcNAcylation itself, accounting for contradictory reports regarding the metabolic requirements of AAM^11–13^. It would be intriguing for further studies to examine how nutrient availability impinges on O-GlcNAcylation to shape IL-4 responsiveness by macrophages. Nevertheless, other studies indicate O-GlcNAcylation is dispensable for IL-4 polarization^19,20^. We hypothesize divergent observations may stem from variable *Ogt* deletion efficiency *in vitro*. Despite this, using multiple models we find O-GlcNAcylation is also required for AAM *in vivo*.

Importantly macrophage O-GlcNAcylation played a key role in systemic immune homeostasis. Mice with macrophage specific *Ogt*-deletion displayed increased T cell activation and IFNγ production. Interestingly, a similar finding was observed using mice with autophagy-deficient macrophages that also lack a stable LCM population. In these mice, the LCM population could be restored by blocking IFNγ signaling^56^. Therefore, cyclic feedback between increased T cell activation, IFNγ production and macrophage death may also contribute in *Lyz2^ΔOgt^* mice. Whether the loss of T cell control occurs mainly because *Lyz2^ΔOgt^* macrophages have reduced suppressive capacity, or is secondary to the absence of a normal tissue macrophage distribution, is uncertain. We and others provide evidence that inhibiting OGT directly alters the ability of macrophages to inhibit T cell activation^57^. Additionally, however, the increase in CD11c^+^ intestinal macrophages signifies more localization towards villus tips^58^, indicating potential increases in microbial interaction^59^. Indeed, during intestinal H.p. infection we observed a marked reduction in LCM and rapid monocyte replacement that is atypical in normal mice. However, this shift in MNP composition in *Lyz2^ΔOgt^* mice highly overlapped with that observed in H.p. infected C57BL/6 mice intraperitoneally co-infected with *S. typhimurium*^60^. Given the minimal separation between intestinal lamina propria and cavity, it is also conceivable that *Lyz2^ΔOgt^* are directly exposed to bacterial ligands in the case of a weakened barrier, particularly during H.p. infection.

Perturbations in macrophage O-GlcNAcylation had further consequences in disease outcomes. Previously, *Lyz2^ΔOgt^* mice exhibited exacerbated high fat diet induced inflammation^19^, but greater resilience to viral infection^23,24^. We additionally show increased susceptibility to helminth infection and better tumor control, of which the latter as also been recently reported^57^. Our data suggest secondary effects on T cell responses and increased IFNγ are responsible, which are known to delay worm expulsion and delay tumor growth^61,62^, rather than direct contributions of the macrophages where alternative activation may be dispensable, at least during H.p. infection^63^. Furthermore, the balance of resident versus newly monocyte-derived macrophages significantly impacts immune responses and resolution, yet this has not previously been accounted for in studies using *Lyz2^ΔOgt^* mice. For instance, monocyte-derived macrophages provide superior anti-viral protection^64^, and thus may account for the resistance of *Lyz2^ΔOgt^* mice to viral infection. Cautious interpretation of the *in vivo* contribution of previously reported mechanisms may therefore be needed.

Together, our work identifies O-GlcNAcylation as a novel regulator of macrophage biology, by governing IL-4 mediated polarization and expansion, as well as controlling the homeostatic turnover of tissue resident populations. This work highlights the potential of O-GlcNAcylation as therapeutic target in diseases with macrophage driven pathology. We show that O-GlcNAcylation is required for macrophage survival during environmental stress, thus in particular its evaluation as a target in scenarios of an altered nutrient microenvironments, such as cardiovascular disease, obesity and tumors, will be highly pertinent.

### Study Limitations

The targets of OGT mediating the observed effects remain to be further studied. Global blockage of O-GlcNAcylation conceivably has considerably different outcomes than single mutations of any of its >10,000 target proteins^65^. For instance, O-GlcNAcylation inhibits activity of glycolytic enzymes such as PFK1^65^, yet promotes glucose metabolism through Myc^66^ and HIF-1α^67^ stabilization. Thus, we focused on the outcome of *Ogt* deletion. The presence of O-GlcNAcylation in *Lyz2^ΔOgt^* macrophage populations hinders identification of truly deficient cells based on staining alone, particularly during models of inflammation in which we observe high turnover of resident macrophage populations. Our data, combined with *Lyz2*-cre reporter mice ubiquitously labelling lung and peritoneal macrophages but not blood monocytes^68,69^, strongly supports a delay between genetic excision and complete loss of protein, or residue removal by OGA, though this in essence provides a tool where progressive loss of O-GlcNAcylation in longer lived resident cells may alleviate negative and potentially pathological effects of a global MNP loss. However, future studies are needed to conclusively determine if the role of O-GlcNAcylation differs between established resident macrophages and differentiating monocytes.

## ACKNOWLEDGEMENTS

Thanks to the funders of this work: Bart Everts is supported by a VIDI grant (91614087) from the Netherlands Organisation for Scientific Research, Rick M Maizels is supported by a Wellcome Trust for my Investigator Award from them (Ref: 219530), and Conor M Finlay is supported by a Research Ireland award (22/PATH-S/10649). We would like to thank the LUMC Flow Core Facility operators for the continual maintenance and trouble-shooting of the instruments, and the animal technicians for their careful considerations and care for animal well-being. Many thanks to our colleagues within the Leiden University Center for Infectious Diseases for continual scientific discussions.

## METHODS

### Animal models

*Lyz2*-cre and *Itgax*-cre mice were crossed with *Ogt^flox^* mice and bred in-house under specific pathogen free (SPF) conditions. Independent experiments were sex and age matched once, but both males and females were used, after reaching a minimum experimental age of 8 weeks. Experiments were performed in accordance with local government regulations, EU Directive 2010/63EU and Recommendation 2007/526/EC regarding the protection of animals used for experimental and other scientific purposes, as well as approved by the Dutch Central Authority for Scientific Procedures on Animals (CCD). Animal license number AVD1160020198846.

#### Peritoneal injections

IL-4 complex was prepared by mixing 5ug of recombinant IL-4 with 25µg of anti-IL4 antibody and injected in 200µl of PBS on days 0 and 2 and mice were sacrificed on the third day. Zymosan was prepared in PBS and mice were injected with 10µg/200µl and sacrificed on day 3. To perform peritoneal cell transfers, total PEC was isolated and pooled for each genotype. *Ogt^flox^* cells were stained with CFSE while *Lyz2^ΔOgt^* mice were stained with CellTrace Violet in sterile PBS for 15 minutes at 37 degrees, washed twice with RPMI/10% FCS, once more with sterile PBS, pooled at ratio of 2:1 *Ogt^flox^*: *Lyz2^ΔOgt^* and resuspended for injection. 1×10^6^ total cells were injected i.p. in 200µl PBS.

#### Infections

*H. polygyrus* infection was given by oral gavage of 200 L3 larvae. For challenge infections were cleared by oral gavage of 2 mg (200µl in ddH_2_O) pyrantel pamoate on day 14 post-infection and effective treatment was confirmed by fecal egg counts. Adult worms counted under a light dissection microscope by gently tearing and rinsing the intestine in warm PBS and removing them with forceps. *S. mansoni* infection was done by percutaneous exposure of 35 cercariae for 30 minutes during anaesthesia with 50mg/Kg ketamine and 10mg/Kg xylazine.

#### Tumor model

B16.F10 melanoma cells were cultured in DMEM containing 10% FCS, 100u/mL penicillin, 100µg/ml streptomycin, 2mM L-glutamine and 1mM pyruvate. Cells were split every 2-3 days and harvested for injection once they reached ~80% confluence. Inoculation was one by injecting 100µl of 3×10^5^ B16.F10 melanoma cells intradermally in the shaved flank during anaesthesia with isoflurane. Mice were monitored 3 times a week for tumor growth, which was calculated by 0.5*length*height*width as measured by a caliper. Sacrifice was performed up to 3 weeks after inoculation or when tumors reached 1500mm^3^.

#### *In vitro* BMDM culture

Femurs and tibias of mice were collected in RPMI, surface sterilized with ethanol and flushed with cold HBSS using a 25 g needle and syringe. Clumps and fragments were removed by aspirating up and down with the syringe and filtering through a 40 μM strainer. Bone marrow cells were centrifuged at 300*g* for 5 min, resuspended and counted in cold RPMI. Cells were adjusted to concentration in RPMI containing 10% heat-inactivated FBS, 100U/ml penicillin, 100 μg/ml streptomycin, 50 μM 2-mercaptoethanol and 10% supernatant of M-CSF producing L929 cells. 2.5 ml containing 2×10^6^ cells were plated in non-culture treated 6-well plates. Cultures were supplemented on day 3 & 5 with 5 ml of complete RPMI containing 20% L929 supernatant. Cells were harvested using accutase on day 7, replated in 24-well plates with 5×10^5^ cells/well and stimulated in 0.5 ml for 24 h with LPS (100 ng/ml) and IFNy (50ng/ml) or IL-4 (40 ng/ml). Plates were put on ice for at least 10 min and BMDM were gently scraped using a 1ml pipet tip to detach cells for downstream assays.

### Gene expression by qPCR

RNA was extracted was performed according to Qiagen RNA easy kit protocol as per manufacturer’s instructions. cDNA synthesis was performed using M-MLV Reverse and qPCR was performed with the Gotaq qPCR master mix and the CFX96 Touch Real-Time PCR Detection System.

### Tissue processing

Cell isolations were done as previously described^32^.

#### Peritoneal exudate cells

5 ml cold PBS containing 2% FBS and 2 mM EDTA was injected into the exposed abdomen, gently agitated for 20 seconds, withdrawn and stored on ice.

#### Intestines

The first 15cm of the small intestine was excised, opened longitudinally over PBS-soaked paper towel. Feces and mucus were gently scraped off with a metal spatula then intestine were washed by vigorous shaking in a 50 ml tube containing 15 ml of Ca/Mg-free HBSS and 2mM EDTA, then cut into 1-2cm pieces and stored in wash buffer on ice until further processing. Three rounds of washing were done in 15ml HBSS/EDTA wash buffer for 20min shaking at 200rpm for 20min. Washed intestines were rinsed once in RPMI before digestion for 15-20 minutes in 10 ml RPMI containing 10% FBS, 1 mg/ml collagenase VIII and 40U/ml DNAse I. Digestion was stopped by adding cold RPMI containing 10% FBS, then resulting suspension was filtered through a 100µM filter, spun at 400g for 5 min, resuspended in PBS/2%FBS/2mM EDTA and filtered through a 40µM filter for counting. Large intestines, including caecum, were processed similarly but digested with 1 mg/ml collagenase IV, 0.5 mg/ml collagenase D, 1 mg/ml dispase II, 40U/ml DNAse I for 25–30 min.

#### Livers

Livers were stored in cold RPMI then minced using a razor blade in a petri plate, and digested similarly to the large intestine with additional manual shaking vigorously by hand every 5–8 min. Digested livers were filtered through 100um strainers, and washed twice with PBS/2%FCS/2mM EDTA, and red-blood cell lysed with ACK lysis buffer. CD45^+^ cells were enriched by positive magnetic selection.

#### Lungs, Spleens, Tumors and Lymph Nodes

Lungs and spleens were stored in cold PBS, minced with scissors and digested in 1 mg/ml collagenase IV and 40U/ml DNAse I and shaken for 30 min before mashing through a 100μm filter. Homogenates were red-blood cell lysed, washed, and passed through a 40μm. Lymph nodes were processed similarly but without red blood cell lysis. Tumor homogenates were subsequently enriched for CD45^+^ cells by magnetic separation.

### Flow cytometry staining and analysis

#### General staining protocol

Cells were plated in a 96-well V-bottom plate, washed with PBS and pre-stained with viability dye and Fc-block in PBS for 15 min at RT. If required, subsequent live surface-staining was performed for fixation sensitive targets in FACS buffer (PBS, 0.5% BSA, 2mM EDTA) for 30 min on ice before fixation. Fixation was performed using eBioscience Foxp3 fixation/permeabilization staining kit for 30-60min at RT. Fixed cells were stored in FACS buffer at 4 degrees until further staining, at which point cells were washed with 1x permeabilization buffer and resuspended in staining mix for 1 hour at RT. Staining mixed contained the remaining antibodies diluted in perm buffer containing 10% brilliant buffer plus and Fc-block. For RELMα, cells were then washed twice and stained with secondary antibody. Stained cells were washed twice with perm buffer and resuspended in FACS buffer for acquisition.

#### Intracellular cytokine staining

For detection of T cell cytokines, total cell homogenates were stimulated with PMA and Ionomycin in the presence of brefeldin A for 4 hours at 37 degrees. Stimulated cells were fixed with 1.8% formaldehyde in PBS for 15 minutes at RT and permeabilized with 1x eBio perm buffer. Staining for surface markers and intracellular cytokines was performed in 1x perm buffer. Stained cells were washed twice and resuspended in FACS for acquisition. Intracellular macrophage cytokines were assessed by culturing the total PEC at 37 degrees in the presence of brefeldin A and monensin for 5 hours. Staining was done according to the general staining protocol.

#### Phospho staining

Cells were fixed with 1.8% formaldehyde in PBS for 20 minutes at RT and permeabilized with 90% methanol in PBS for 20 minutes at 4 degrees. Cells were washed twice in PBS and stained for phospho targets for either 1 hour at RT with additional surface antibodies, or overnight at 4 degrees for pH2AX and pSTAT6 and subsequently stained for surface antibodies the next day.

#### Metabolic dyes

Single-cell suspensions were plated in a non-treated V-bottom 96 well plate, washed with pre-warmed RPMI and suspended in 100ul of un-supplemented RPMI containing 5 nM TMRM and 20nM MitoTracker DeepRed, or 1µM DCFDA for 20 min at 37 °C. Cells were washed twice in FACS buffer before lived/dead and surface staining. Cells were acquired immediately after staining in FACS buffer.

#### Analysis

All acquisition was performed on Cytek 5-laser spectral analyzers. Data was unmixed using Cytek SpectroFlo software version 3 as outline previously (ref). Unmixed data was analysis with FlowJo version 9 and additional visualized with GraphPad Prism version 8 or R Studio version 4.

### Electron Microscopy

#### Fixing, embedding and acquisition

Total PEC was isolated in 3ml PBS and directly fixed by adding 2X concentrated fixation buffer ultimately containing a final concentration of 1.5% glutaraldehyde and 0.1M cacodylate. Cells were centrifuged at 2500rpm for 10 minutes, rinsed with 0.1M cacodylate buffer, and placed in osmium tetroxide (1% OsO4 and 1.5% potassium hexacyanoferrate (III) in MilliQ), and left on ice for 1 hour. Fixed cells were centrifuged at 4000rpm for 5 minutes and rinsed with 0.1M cacodylate buffer, then resuspended in agar (2% bacto-agar in MilliQ). Samples were again centrifuged at 5000 rpm for 5 minutes. Once the agar had set, the pellet was cut off and stored in 70% ethanol overnight at 4°C. Cells were subsequently dehydrated using a series of 80% and 90% ethanol baths for 10 minutes each, before being put in 100% absolute ethanol twice for 30 minutes. To embed the fixed cells, they were bathed in subsequent Epon/Acetone ratios of 1:2, 1:1, and 2:1 each for 30 minutes, followed by an hour of 100% Epon. Fully embedded samples were transferred to BEEM® capsules and left for two days at 70°C. Hardened samples were sliced 90 nm or 45 nm thick using a diamond knife on an ultra-microtome (Leica EM UC6) and captured on a copper grid (mesh 50). Copper grids were exposed to a droplet of 7% uranyl acetate for 10 minutes to stain the cells, then washed with 5 droplets of MilliQ and treated with 0.01M NaOH. Finally, a droplet of lead citrate was added to the grid for 5 minutes, which was subsequently washed 6 times using 0.01M NaOH. Air-dried grids were then ready for imaging. Images were acquired using an Tecnai 12 Spirit Twin transmission electron microscope (FEI) equipped with an Eagle 4k x 4k CCD camera (FEI). Images were taken at 6500x nominal magnification at 120 kV. An emission current between 0.5 µA and 2.5 μA and a defocus of 2 µm was used for all images. MyTEM and MyStitch software 4 was used to record consecutive images with a 20% overlap and stitch these into a composite virtual slide. Images were saved using Aperio ImageScope (Leica).

#### Annotation and supervised Machine Learning

Cell and organelle annotations were done manually prior to machine learning application. Macrophages were distinguished by cell size and the presence of endocytic vesicles, that are absent in lymphocytes. Nuclei were easily distinguished, in which dark areas were defined as heterochromatin, and lighter areas marked as euchromatin. Mitochondria were defined by having a double membrane and cristae. For machine learning, stitches were processed in CAVIA (Koning et al. Manuscript in preparation). Virtual boxes containing manually annotated structures of interest were placed to define the regions used for network training. Models were created using the TensorFlow and Keras platforms 13. Typically, three rounds of annotation, modelling, and predictions (initial annotation, removal of false positives and addition of false negatives) were used for model optimization. Minor manual corrections on the final predictions of the cell and nucleus were done prior to data quantification.

#### Image analysis and quantification

Segmentations of predicted or manually annotated structures were extracted with a custom Python 3.8.10 script using Spyder 5.5.3. Occurrence, size, and fill factor were calculated. To calculate fill factor either whole macrophages, or nuclei were selected as a primary segmentation, in which the presence of a secondary segmentation (nucleus, mitochondria for macrophages; heterochromatin and euchromatin for nuclei) was calculated. Occurrence represented the number of secondary structures within the primary one, size was calculated as the total area of the segmented objects, and fill factor represents the relative total area compared to the primary structure. All predictions of the macrophages and nuclei were manually corrected to optimize accuracy.

### Confocal Microscopy

Total PEC or BMDM were plated on Ibidi 8 well chamber coverslips in serum-free media for 1 hour, washed with warm PBS 3 times with gently agitation and fixed with either 2% ultrapure formaldehyde for 15 minutes at RT and stained with surface markers and DAPI for 30 minutes at RT, followed by LipidTox staining 1 hour before acquisition, or with eBioscience fixation buffer and stained for intracellular targets 1 hour at RT in perm buffer, and subsequently DAPI for 10 min in PBS. Acquisition done with a Leica SP8X white light laser microscope and processed using the Leica LASX software.

### Single-cell RNA sequencing

#### Sample preparation and sequencing

80.000 total PEC cells were plated in separate wells of a V-bottom plate and each sample stained on ice with 1ul of an individual TotalSeqB hashtag in 50ul of PBS. Cells were washed three times with PBS and pooled a concentration of 2000 cell/ul in PBS containing 0.04% BSA for sequencing. Cells had greater than 95% viability before sample submission, and a target of 3000 cells/sample were sequenced (~18,000 from pooled sample). scRNA-seq libraries were obtained using the Chromium Next GEM Single Cell 3’ Library & Gel Bead Kit v3.1 (10x Genomics) and Chromium Next GEM Chip G Single Cell Kit (10x Genomics) following the manufacturer’s protocol (CG000317_RevD). Libraries were sequenced on an Illumina NovaSeq 6000 and reads were mapped and counted with Cell Ranger v7.0.0.

#### Analysis

Cellranger identified a total of 14,188 cells. Downstream analysis was performed in the Seurat and Bioconductor environment in R (v.4.3.2-4.4.2). Sample demultiplexing and inter-sample doublet identification was identified using the HTODemux() function, identifying 12,045 singlets. Cells were filtered using quality control metrics, removing cells with >20% mitochondrial reads, >60,000 total RNA UMIs, or expressing fewer than 100 or more than 7,500 unique genes. Countd data using the Seurat function NormalizeData(), genes with high variance were identified using the FindVariableFeatures() function, with the “vst” method, selecting 2,000 features for downstream analysis. These genes were scaled using the ScaleData() function and used to produce a principle component analysis (PCA). The top 30 PCs were used to produce a UMAP using the RunUMAP() function. A graph based (Louvain) clustering was performed on the first 30 PCs using the FindNeighbours() and FindClusters() with resolution = 0.2 deliberately under-clustering to return only major cell lineages. Cell populations were manually annotated based on canonical marker genes and included B1 cells, B2 cells, macrophages, ILC/T lymphocytes, dendritic cells, and mast cells. Cell cycle scoring was performed using the CellCycleScoring() function with a curated list of S and G2M phase markers converted from human to mouse gene nomenclature.

#### Pyscenic

Single cell gene regulatory network analysis was calculated using PySCENIC (Reference) v0.1.12 using normalized gene expression input. Cells were pre-filtered to match quality control cutoffs. The analysis was performed on the full dataset after cell quality control filtering and on the smaller MNP dataset using all the version 9 (mc9nr) cisTARGET databases (https://resources.aertslab.org/cistarget/).

FeatherRankingDatabase(name=“mm9-tss-centered-10kb-10species.mc9nr.genes_vs_motifs.rankings”), FeatherRankingDatabase(name=“mm9-tss-centered-5kb-10species.mc9nr.genes_vs_motifs.rankings”), FeatherRankingDatabase(name=“mm9-500bp-upstream-10species.mc9nr.genes_vs_motifs.rankings”), FeatherRankingDatabase(name=“mm9-tss-centered-5kb-7species.mc9nr.genes_vs_motifs.rankings”), FeatherRankingDatabase(name=“mm9-tss-centered-10kb-7species.mc9nr.genes_vs_motifs.rankings”), FeatherRankingDatabase(name=“mm9-500bp-upstream-7species.mc9nr.genes_vs_motifs.rankings”)

SCENIC regulon activity scores were integrated with Seurat RNA object as a new assay. The AUC matrix from SCENIC was scaled and PCA was performed on the regulon data from which a UMAP was generated using the top 20 PCs. Graph-based clustering was performed on the regulon principal components using FindNeighbors() and FindClusters() functions (resolution = 0.25). Additionally, hierarchical clustering was performed on the regulon principal component space using Ward’s method with Euclidean distances, and dynamic tree cutting was applied to define cluster relationships (cutreeDynamic, minimum cluster size = 5). The regulon-based clustering validated the major cell populations identified through transcriptional analysis.

#### Macrophage subcluster analysis

Subsequent analysis focused on the MNP compartment following initial lineage identification. Additional quality control was performed on the MNP subset, removing cells with ribosomal gene content <10% or mitochondrial content >8% to exclude low-quality cells while retaining biological heterogeneity. The filtered dataset underwent standard normalization and scaling, with 1,000 variable features selected for downstream analysis. PCA was performed, and the first 30 PCs were used for UMAP visualization and graph-based clustering (resolution = 1.0). Clusters were manually annotated based on canonical marker expression including Zbtb46, Itgax (dendritic cells), Sema4a, Cd74, H2-Aa (small cavity macrophages), Mrc1, Trem2, Cd163 (converting cavity macrophages), and Gata6, Nt5e, Tgfb2 (large cavity macrophages). Cell cycle states were determined using mouse orthologs of established S and G2M phase markers. This analysis identified eleven distinct populations: cDC1, cDC2, DC-like small cavity macrophages (SCM), monocyte-like SCM, converting cavity macrophages (CCM), altered large cavity macrophages (LCM), LCM, and cells in S and G2M phases. These populations were further validated through both gene expression and regulon activity analysis. Weighted nearest neighbor analysis was performed integrating both transcriptional and regulon data to generate a unified representation of cellular states. Final clustering was performed on this integrated representation using the Louvain algorithm (resolution = 0.5), confirming the stability of the identified populations.

SCM-LCM scores were constructed from Immunological Genome Project datasets (version 1 microarray) ‘MF.II-480hi.PC’ for LPM and ‘MF.II+480lo.PC’ for SCM. Pseudo-time trajectory was generated using TSCAN (Tools For Single-Cell Analysis) after subsetting *Ogt^flox^* cells and excluding DC/DC-like SCM. Seurat_cluster 3, which spanned monocyte-like SCM and CCM was split according to the corresponding MNP identity and the cluster 3 SCM were used as the start of the trajectory. RNA Velocity was performed on a spliced unspliced calculation for each transcript. To harmonize across analysis pipelines we filtered the slicing matrix to retain only those genes and cells present after QC steps in our R analysis pipeline and converting our seurat object to an AnnData object, and thus inheriting clustering information from R. Due to minor differences in normalization and calculation of variable genes we performed the core scanpy analysis pipeline to rederive a PCA and used the top 50 PC tom compute a UMAP. RNA velocity analysis was performed with the scvelo package using the top 2000 genes and a miniumn of 10 shared counts in the scv.pp.filter_and_normalize() function. The top 50 PCs and nearest neighbors = 50 for sc.pp.moments() function. For RNA velocity calculations we used scv.tl.recover_dynamics and scv.tl.velocity functions using the dynamical model.

### Metabolic Measurements

#### Seahorse

Peritoneal macrophages were sorted directly in 96 well plates at 80.000 cells/well using a Beckman Coulter CytoFlex SRT cell sorter. Sorted cells were washed in Seahorse XF assay media and transferred to a Seahorse XF assay plate in 80ul of assay media containing 10mM glucose, 2mM Glutamax and 2mM sodium-pyruvate. Plated cells were spun at low g to form a monolayer and incubated for 1 hour at 37 degrees in a non-CO_2_ incubator. Prior to analysis 95µl of fresh assay media was added to each well. Pre-hydrated cartridges were prepared during cell incubation. Mito-stress assay was performed using 2 μM oligomycin, 1.5 μM FCCP, and 0.75 μM each of rotenone and antimycin A.

#### Click-chemistry and ATP dependencies

HPG uptake and analysis done as previously^32,47^. 5×10^5^ of total PEC was plated in a 96-well V-bottom plate in 90ul of methionine-free media containing 65 mg/L l-cystine dihydrochloride, 1x glutamax and 10% dialyzed FCS. Cells were methionine starved for 45 min before addition of either 2 µM oligomycin, 100 mM 2DG, or both for 15 min. A final concentration of 100 µM Homopropargylglycine (HPG) was added for 30 min before performing staining an click reaction. After live/dead staining, fixation was done with 2% ultrapure formaldehyde and permeabilized with 1% BSA/0.1% saponin in PBS for 15 min. Permeabilized cells were washed twice with PBS and stained with the click reaction mix made by sequential addition of 100 mM Sodium Ascorbate (10mM final), 200 mM THPTA (2mM final), and 1mM AFdye488 azide plus (0.5 µM final) to 50mM CuSO_4_ (0.5 mM final conc.) and made up to volume with PBS. Samples were incubated for 30 min in the dark at room temperature. Cells were acquired in FACS buffer after surface staining. Metabolic dependencies were calculated as shown previously^48^.

### Western blotting

Proteins were collected from by lysis in 100 µl RIPA buffer supplemented with 1x protease inhibitor, 10 µM OGA inhibitor Thiamet-G and 20 µM OGT inhibitor ST045849 to prevent removal of the post-translational modifications. After 30 min on ice, lysate was centrifuged at maximum speed for 5 min at 4 degrees to get rid of cell debris. Protein concentration in supernatant was quantified using the Pierce BCA protein assay kit and equal amounts of proteins were boiled at 95° for 5 min. Proteins were separated by SDS-PAGE and transferred to PVDF membranes using semi-dry method with Trans-Blot Turbo Transfer system. Membranes were blocked for 1 hour at RT in TTBS buffer (20 mM Tris–HCl [pH 7.6], 137 mM NaCl, and 0.25% [v/v] Tween 20) containing 5% milk and incubated overnight at 4 degrees with primary antibodies diluted 1:1000 in TTBS containing 5% BSA. The next day, membranes were washed in TTBS and incubated for 2 hours at RT with HRP-conjugated secondary antibodies. After washing, signal developed using Clarity Western ECL substrate and quantified using ImageJ. Signal from protein of interest was then normalized to housekeeping gene and reference condition.

### Histology

Liver sections were placed in 3.7% formaldehyde/PBS overnight at 4 degrees, transferred to cassettes and stored in 70% ethanol until paraffin embedding could be performed. Slides were prepared using 5µM sections cut using a rotary microtome and submitted to LUMC Pathology Department for H & E staining. Images were captured using a 3DHISTECH Pannoramic 250 digital scanner. 20X and 10X magnification snapshots were taken for determining granuloma size and counts respectively, which were subsequently quantified in ImageJ by manually circling granulomas and taking the average counts and areas from 5 fields/sample.

### Statistical Analysis

GraphPad Prism was used for statistical testing. Unless stated otherwise stated, summary lines and bars in plots represent the mean, with individual biological replicates shown. Test details are present within the figure legends. Where p values are not numerically stated, annotated symbols represent: *p < 0.05, **p < 0.01, ***p < 0.001, ****p < 0.0001 and ns = not significant.

## Notes

### Competing Interest Statement

The authors have declared no competing interest.

## REFERENCES

1. Tannahill, G. et al. Succinate is a danger signal that induces IL-1β via HIF-1α. Nature 496, 238–242 (2013).

2. Verberk, S. G. S. et al. An integrated toolbox to profile macrophage immunometabolism. Cell Rep Methods 2, 100192 (2022).

3. Baardman, J. et al. A Defective Pentose Phosphate Pathway Reduces Inflammatory Macrophage Responses during Hypercholesterolemia. Cell Reports 25, 2044–2052.e5 (2018).

4. Castoldi, A. et al. Triacylglycerol synthesis enhances macrophage inflammatory function. Nat Commun 11, 4107 (2020).

5. Rosas-Ballina, M., Guan, X. L., Schmidt, A. & Bumann, D. Classical Activation of Macrophages Leads to Lipid Droplet Formation Without de novo Fatty Acid Synthesis. Front Immunol 11, 131 (2020).

6. Wu, K. K. et al. MDM2 induces pro-inflammatory and glycolytic responses in M1 macrophages by integrating iNOS-nitric oxide and HIF-1α pathways in mice. Nat Commun 15, 8624 (2024).

7. Huang, S. C.-C. et al. Metabolic Reprogramming Mediated by the mTORC2-IRF4 Signaling Axis Is Essential for Macrophage Alternative Activation. Immunity 45, 817–830 (2016).

8. Liu, P.-S. et al. α-ketoglutarate orchestrates macrophage activation through metabolic and epigenetic reprogramming. Nat Immunol 18, 985–994 (2017).

9. Vats, D. et al. Oxidative metabolism and PGC-1β attenuate macrophage-mediated inflammation. Cell Metab 4, 13–24 (2006).

10. Huang, S. C.-C. et al. Cell-intrinsic lysosomal lipolysis is essential for alternative activation of macrophages. Nat Immunol 15, 846–855 (2014).

11. Nomura, M. et al. Fatty acid oxidation in macrophage polarization. Nat Immunol 17, 216–217 (2016).

12. Wang, F. et al. Glycolytic Stimulation Is Not a Requirement for M2 Macrophage Differentiation. Cell Metab 28, 463–475.e4 (2018).

13. Divakaruni, A. S. et al. Etomoxir Inhibits Macrophage Polarization by Disrupting CoA Homeostasis. Cell Metab 28, 490–503.e7 (2018).

14. Namgaladze, D. & Brüne, B. Fatty acid oxidation is dispensable for human macrophage IL-4-induced polarization. Biochim Biophys Acta 1841, 1329–1335 (2014).

15. Ramakrishnan, P. O-GlcNAcylation and immune cell signaling: A review of known and a preview of unknown. J Biol Chem 300, 107349 (2024).

16. Jha, A. K. et al. Network integration of parallel metabolic and transcriptional data reveals metabolic modules that regulate macrophage polarization. Immunity 42, 419–430 (2015).

17. Hinshaw, D. C. et al. Hedgehog Signaling Regulates Metabolism and Polarization of Mammary Tumor-Associated Macrophages. Cancer Res 81, 5425–5437 (2021).

18. Zhao, M. et al. Epithelial STAT6 O-GlcNAcylation drives a concerted anti-helminth alarmin response dependent on tuft cell hyperplasia and Gasdermin C. Immunity 55, 623–638.e5 (2022).

19. Yang, Y. et al. OGT suppresses S6K1-mediated macrophage inflammation and metabolic disturbance. Proc Natl Acad Sci U S A 117, 16616–16625 (2020).

20. Shi, Q. et al. Increased glucose metabolism in TAMs fuels O-GlcNAcylation of lysosomal Cathepsin B to promote cancer metastasis and chemoresistance. Cancer Cell 40, 1207–1222.e10 (2022).

21. Li, X. et al. O-GlcNAc transferase suppresses inflammation and necroptosis by targeting receptor-interacting serine/threonine-protein kinase 3. Immunity 50, 576–590.e6 (2019).

22. Al-Mukh, H. et al. Lipopolysaccharide Induces GFAT2 Expression to Promote O-Linked β-N-Acetylglucosaminylation and Attenuate Inflammation in Macrophages. J Immunol 205, 2499–2510 (2020).

23. Wang, Q. et al. O-GlcNAc transferase promotes influenza A virus–induced cytokine storm by targeting interferon regulatory factor–5. Sci Adv 6, eaaz7086 (2020).

24. Li, T. et al. O-GlcNAc transferase links glucose metabolism to MAVS-mediated antiviral innate immunity. Cell Host Microbe 24, 791–803.e6 (2018).

25. Song, N. et al. MAVS O-GlcNAcylation Is Essential for Host Antiviral Immunity against Lethal RNA Viruses. Cell Rep 28, 2386–2396.e5 (2019).

26. Mass, E., Nimmerjahn, F., Kierdorf, K. & Schlitzer, A. Tissue-specific macrophages: how they develop and choreograph tissue biology. Nat Rev Immunol 23, 563–579 (2023).

27. Li, W., Yang, Y., Yang, L., Chang, N. & Li, L. Monocyte-derived Kupffer cells dominate in the Kupffer cell pool during liver injury. Cell Rep 42, 113164 (2023).

28. Bain, C. C., Oliphant, C. J., Thomson, C. A., Kullberg, M. C. & Mowat, A. M. Proinflammatory Role of Monocyte-Derived CX3CR1int Macrophages in Helicobacter hepaticus-Induced Colitis. Infect Immun 86, e00579–17 (2018).

29. Louwe, P. A. et al. Recruited macrophages that colonize the post-inflammatory peritoneal niche convert into functionally divergent resident cells. Nat Commun 12, 1770 (2021).

30. Aegerter, H. et al. Influenza-induced monocyte-derived alveolar macrophages confer prolonged antibacterial protection. Nat Immunol 21, 145–157 (2020).

31. Oh, M.-H. et al. mTORC2 signaling selectively regulates generation and function of tissue-resident peritoneal macrophages. Cell Rep 20, 2439–2454 (2017).

32. Heieis, G. A. et al. Metabolic heterogeneity of tissue-resident macrophages in homeostasis and during helminth infection. Nat Commun 14, 5627 (2023).

33. Gautier, E. L. et al. Gata6 regulates aspartoacylase expression in resident peritoneal macrophages and controls their survival. J Exp Med 211, 1525–1531 (2014).

34. Ong, Q., Han, W. & Yang, X. O-GlcNAc as an Integrator of Signaling Pathways. Front. Endocrinol. 9, (2018).

35. Saunders, H., Dias, W. B. & Slawson, C. Growing and dividing: how O-GlcNAcylation leads the way. J Biol Chem 299, 105330 (2023).

36. Finlay, C. M. et al. T helper 2 cells control monocyte to tissue-resident macrophage differentiation during nematode infection of the pleural cavity. Immunity 56, 1064–1081.e10 (2023).

37. Bain, C. C. et al. Long-lived self-renewing bone marrow-derived macrophages displace embryo-derived cells to inhabit adult serous cavities. Nat Commun 7, ncomms11852 (2016).

38. Bain, C. C., et al. Rate-of-replenishment and microenvironment contribute to the sexually dimorphic phenotype and function of peritoneal macrophages. Sci Immunol 5, eabc4466 (2020).

39. Pesce, J. T., et al. Arginase-1–Expressing Macrophages Suppress Th2 Cytokine–Driven Inflammation and Fibrosis. PLOS Pathogens 5, e1000371 (2009).

40. Huber, S., Hoffmann, R., Muskens, F. & Voehringer, D. Alternatively activated macrophages inhibit T-cell proliferation by Stat6-dependent expression of PD-L2. Blood 116, 3311–3320 (2010).

41. Jarjour, N. N. et al. Bhlhe40 mediates tissue-specific control of macrophage proliferation in homeostasis and type 2 immunity. Nat Immunol 20, 687–700 (2019).

42. Saul, D. et al. A new gene set identifies senescent cells and predicts senescence-associated pathways across tissues. Nat Commun 13, 4827 (2022).

43. Härtlova, A. et al. DNA Damage Primes the Type I Interferon System via the Cytosolic DNA Sensor STING to Promote Anti-Microbial Innate Immunity. Immunity 42, 332–343 (2015).

44. Yu, Q. et al. DNA-Damage-Induced Type I Interferon Promotes Senescence and Inhibits Stem Cell Function. Cell Reports 11, 785–797 (2015).

45. Abbadie, C. & Pluquet, O. Unfolded Protein Response (UPR) Controls Major Senescence Hallmarks. Trends in Biochemical Sciences 45, 371–374 (2020).

46. Ogrodnik, M. et al. Guidelines for minimal information on cellular senescence experimentation in vivo. Cell 187, 4150–4175 (2024).

47. Vrieling, F. et al. CENCAT enables immunometabolic profiling by measuring protein synthesis via bioorthogonal noncanonical amino acid tagging. Cell Rep Methods 4, 100883 (2024).

48. Argüello, R. J. et al. SCENITH: A flow cytometry based method to functionally profile energy metabolism with single cell resolution. Cell Metab 32, 1063–1075.e7 (2020).

49. Bain, C. C. et al. Resident and pro-inflammatory macrophages in the colon represent alternative context-dependent fates of the same Ly6Chi monocyte precursors. Mucosal Immunol 6, 498–510 (2013).

50. Vanneste, D. et al. MafB-restricted local monocyte proliferation precedes lung interstitial macrophage differentiation. Nat Immunol 24, 827–840 (2023).

51. Zhu, L. et al. TSC1 controls macrophage polarization to prevent inflammatory disease. Nat Commun 5, 4696 (2014).

52. Weichhart, T. mTOR as regulator of lifespan, aging and cellular senescence. Gerontology 64, 127–134 (2018).

53. Chen, J. et al. Inhibition of O-GlcNAc transferase activates type I interferon-dependent antitumor immunity by bridging cGAS-STING pathway. eLife 13, RP94849 (2024).

54. Chen, Q. & Yu, X. OGT restrains the expansion of DNA damage signaling. Nucleic Acids Res 44, 9266–9278 (2016).

55. Zhou, Z. et al. Type 2 cytokine signaling in macrophages protects from cellular senescence and organismal aging. Immunity 57, 513–527.e6 (2024).

56. Wang, Y.-T. et al. Select autophagy genes maintain quiescence of tissue-resident macrophages and increase susceptibility to Listeria monocytogenes. Nat Microbiol 5, 272–281 (2020).

57. Zhang, Y. et al. EGR2 O-GlcNAcylation orchestrates the development of protumoral macrophages to limit CD8+ T cell antitumor responses. Cell Chem Biol 32, 809–825.e7 (2025).

58. Corbin, A. L., et al. IRF5 guides monocytes towards an inflammatory CD11c+ macrophage phenotype and promotes intestinal inflammation. Sci Immunol 5, eaax6085 (2020).

59. Kang, B. et al. Commensal microbiota drive the functional diversification of colon macrophages. Mucosal Immunol 13, 216–229 (2020).

60. Rückerl, D. et al. Macrophage origin limits functional plasticity in helminth-bacterial co-infection. PLoS Pathog 13, e1006233 (2017).

61. Salerno, F. et al. Critical role of post-transcriptional regulation for IFN-γ in tumor-infiltrating T cells. Oncoimmunology 8, e1532762 (2018).

62. Kapse, B. et al. Age-dependent rise in IFN-γ competence undermines effective type 2 responses to nematode infection. Mucosal Immunol 15, 1270–1282 (2022).

63. Westermann, S. et al. Th2-dependent STAT6-regulated genes in intestinal epithelial cells mediate larval trapping during secondary Heligmosomoides polygyrus bakeri infection. PLoS Pathog 19, e1011296 (2023).

64. Li, F., et al. Monocyte-derived alveolar macrophages autonomously determine severe outcome of respiratory viral infection. Sci Immunol 7, eabj5761 (2022).

65. Yi, W. et al. Phosphofructokinase 1 glycosylation regulates cell growth and metabolism. Science 337, 975–980 (2012).

66. Swamy, M. et al. Glucose and glutamine fuel protein O-GlcNAcylation to control T cell self-renewal and malignancy. Nat Immunol 17, 712–720 (2016).

67. Ferrer, C. M. et al. O-GlcNAcylation regulates cancer metabolism and survival stress signaling via regulation of HIF-1 pathway. Mol Cell 54, 820–831 (2014).

68. McCubbrey, A. L., Allison, K. C., Lee-Sherick, A. B., Jakubzick, C. V. & Janssen, W. J. Promoter Specificity and Efficacy in Conditional and Inducible Transgenic Targeting of Lung Macrophages. Front. Immunol. 8, (2017).

69. Abram, C. L., Roberge, G. L., Hu, Y. & Lowell, C. A. Comparative analysis of the efficiency and specificity of myeloid-Cre deleting strains using ROSA-EYFP reporter mice. J Immunol Methods 408, 89–100 (2014).

