## Supplemental Figures for "O-GlcNAcylation drives macrophage IL-4 responsiveness and tissue residency through metabolic and cell cycle calibration"

#### SUPPLEMENARY FIGURE LEGENDS:

##### Figure S1. Immune phenotyping of Lyz2Ogt mice during helminth infection

(**A & B**) Gating strategies for the small intestine (A) and PEC (B).

(**C**) Representative cytokine staining following PMA/ionomycin stimulation and quantification of IL-17, Foxp3+ Tregs and IFNγ across tissues during 1° H.p. infection. Pooled from two or three experiments, n=2-7 mice/group. Statistics calculated using two-way ANOVA with Sidak correction for mean row comparisons.

(**D** & **E**) Quantification of (D) DC and DC subset frequency, and (E) expression of activation markers at day 7 of 1° H.p. infection. Representative of two experiments, n=3-6.

(**F** & **G**) Frequency (F) and activation marker expression (G) of splenic DC and their subsets. Representative of two experiments.

##### Figure S2. OGT is required for alternative macrophage activation in multiple models

(**A**) Average egg granuloma count and surface area from the 16 week S.m. infected liver. Data from one experiment with n=8-11/group.

(**B**) Disappearance of $TIM4^{+}$ liver resident macrophage during chronic S.m. infection.

(**C**) Gating of CD45+ cells for differentiating and mature macrophages and evaluation of alternative activation markers during liver infection.

(**D** & **E**) Characterization of (D) $CD4^{+}$ and (E) $CD8^{+}$ T cell phenotypes during S.m. infection.

(**F**) Mean and individual tumor growth curves, and end weight, pooled from two experiments, n = 7-12/group.

(**G**) Gating and quantification for tumor macrophage populations.

(**H**) Dimensionality reduction of intra-tumoral CD4 and CD8 T cells. Targets: TIM3, PD1, CD44, CD62L, CD27, CD3, CD25, CD69, KLRG1.

(**I**) IFNγ expression of CD8 T cells from the draining and non-draining lymph node.

Statistics calculated using (C, G & I) unpaired two-tailed t-test with Welch's correction in cases of unequal variance, (D) two-way ANOVA with Sidak correction for mean row comparisons and (F) Mixed effects model with Sidak multiple comparisons test.

##### Figure S3. Identification of MNP populations using scRNA-seq

(**A**) Representative peritoneal MNP populations and characteristics from mice used for scRNA-sequencing.

(**B**) Number of cells per sample after hashtag oligo (HTO) de-multiplexing.

(**C**) Quality control filters for downstream analysis.

(**D-F**) Identification of macrophages in RNA-UMAP space from 12,045 filtered cells, grouped by genotype or sample.

(**G**) Identification of macrophage cluster using gene regulatory analysis via SCENIC.

(**F**) Removal of Macrophage and B cell doublets.

(**I & J**) Subset macrophage cluster UMAP

(**K & L**) Quantification of lineage clusters and MNP subsets.

(**M-O**) Identification of initial 11 MNP sub-clusters and subsequent manual annotation and merging to define 8 final populations used for analysis.

#### Figure S4. BHLHE40 is not O-GlcNAcylated despite overlap between OGT and BHLHE40 deficiency

(**A**) MNP populations scored against similarity to microarray dataset of BHLHE40 knockout peritoneal macrophages, overlayed on UMAP space and ranked according to score.

(**B**) Confocal microscopy of BHLHE40 expression and O-GlcNAcylation, and quantification of total BHLHE40 signal in O-GlcNAc$^{+}$ or O-GlcNAc$^{-}$ peritoneal macrophages.

(**C**) Western blot for total BHLHE40 or O-GlcNAc after BHLHE40 pull-down using lysate of GM-CSF differentiation BMDM.

(**D**) Pull-down and western blot analysis for BHLHE40 and O-GlcNAc using Alveolar macrophage-like BMDM, differentiated with GM-CSF, TGF-β and PPARγ agonist.

#### Figure S5. OGT regulates macrophage metabolism in vivo and in vitro

(**A**) Violin plots for fatty acid binding proteins, perilipin 1 and lipoprotein lipase, with corresponding feature plots.

(**B**) LipidTox DeepRed staining of unstimulated BMDM. Statistics done using unpaired t-test.

(**C**) Representative histograms for metabolic dye staining of wild-type BMDM after overnight treatment with OGTi.

(**D**) Representative confocal images of M-CSF BMDM, and correspond flow cytometry dot plots for forward and side scatter.

(**E**) Metabolic target expression determined by flow cytometry in unstimulated BMDM, with heatmap showing fold change from control *flox* BMDM.

(**F**) Representative cell-cycle analysis of BMDM.

#### Figure S6. Macrophage residency and expansion requires dynamic O-GlcNAcylation across tissues

(**A**) Number of lung or liver macrophages and change in O-GlcNAc staining after in vivo IL-4c treatment. Representative of two experiments.

(**B**) Feature and violin plots of *Ogt* and *Oga* expression from scRNA-seq data of peritoneal macrophages.

(**C**) Frequency and number of tissue macrophages from Itgax-cre Ogtflox mice. Pooled from 2-3 experiments, n = 3-5 mice/group.

Statistics performed using two-tailed unpaired t-test or two-way ANOVA with Sidak correction for mean row comparisons.

##### Figure S7. OGT controls macrophage residency independent of sex

(**A**) Retrospective pooling and analysis of sex or age differences in peritoneal macrophage composition. Pooled from 11 experiments.

(**B**) Retrospective pooling and analysis of indicated tissue macrophage populations according to sex. Pooled from 5-7 experiments.

Statistics performed using two-way ANOVA with Sidak correction for mean row comparisons.

### Figure S1 - Related to main Figure 2

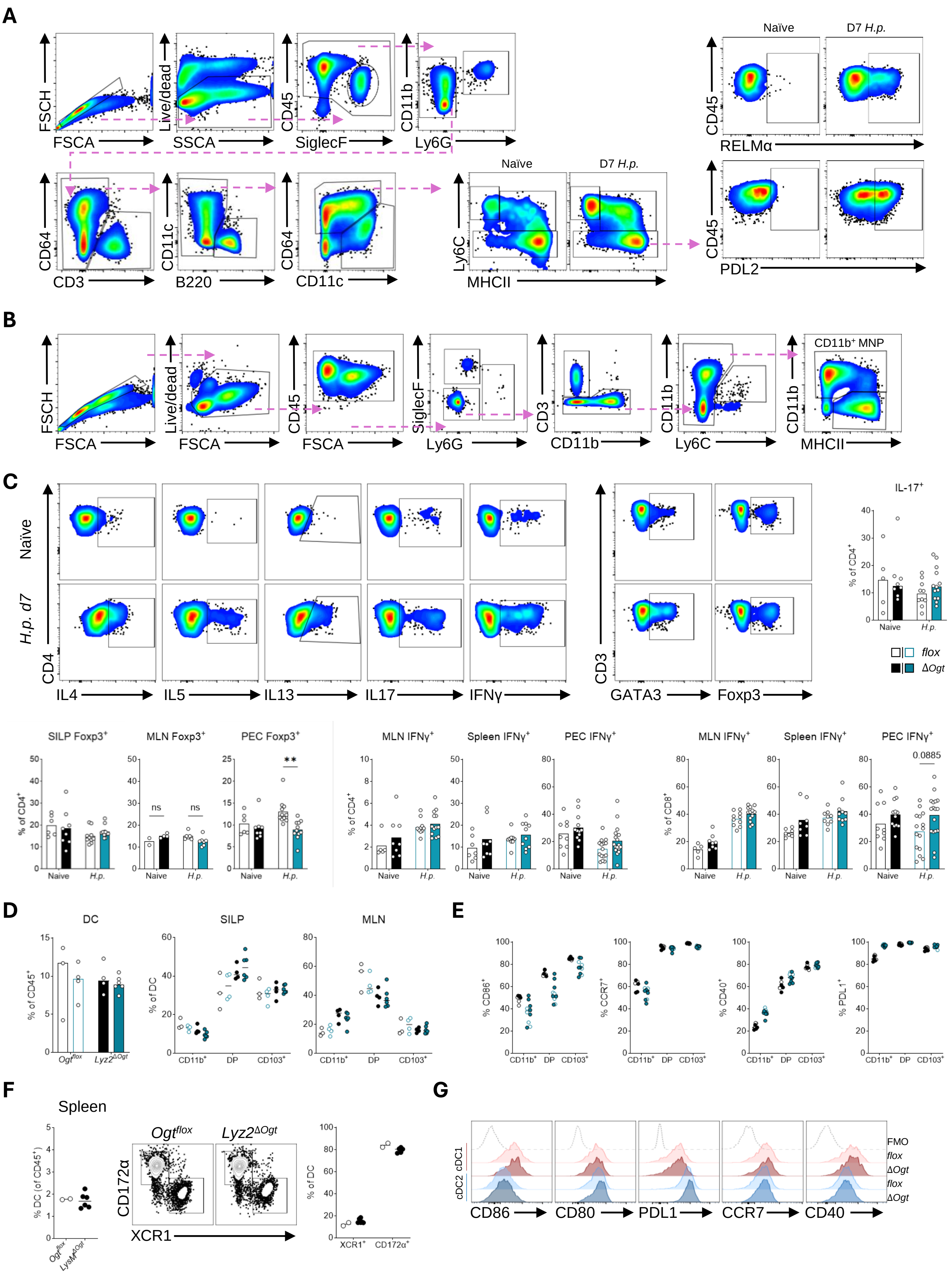

### Figure S2 – Related to main Figure 2

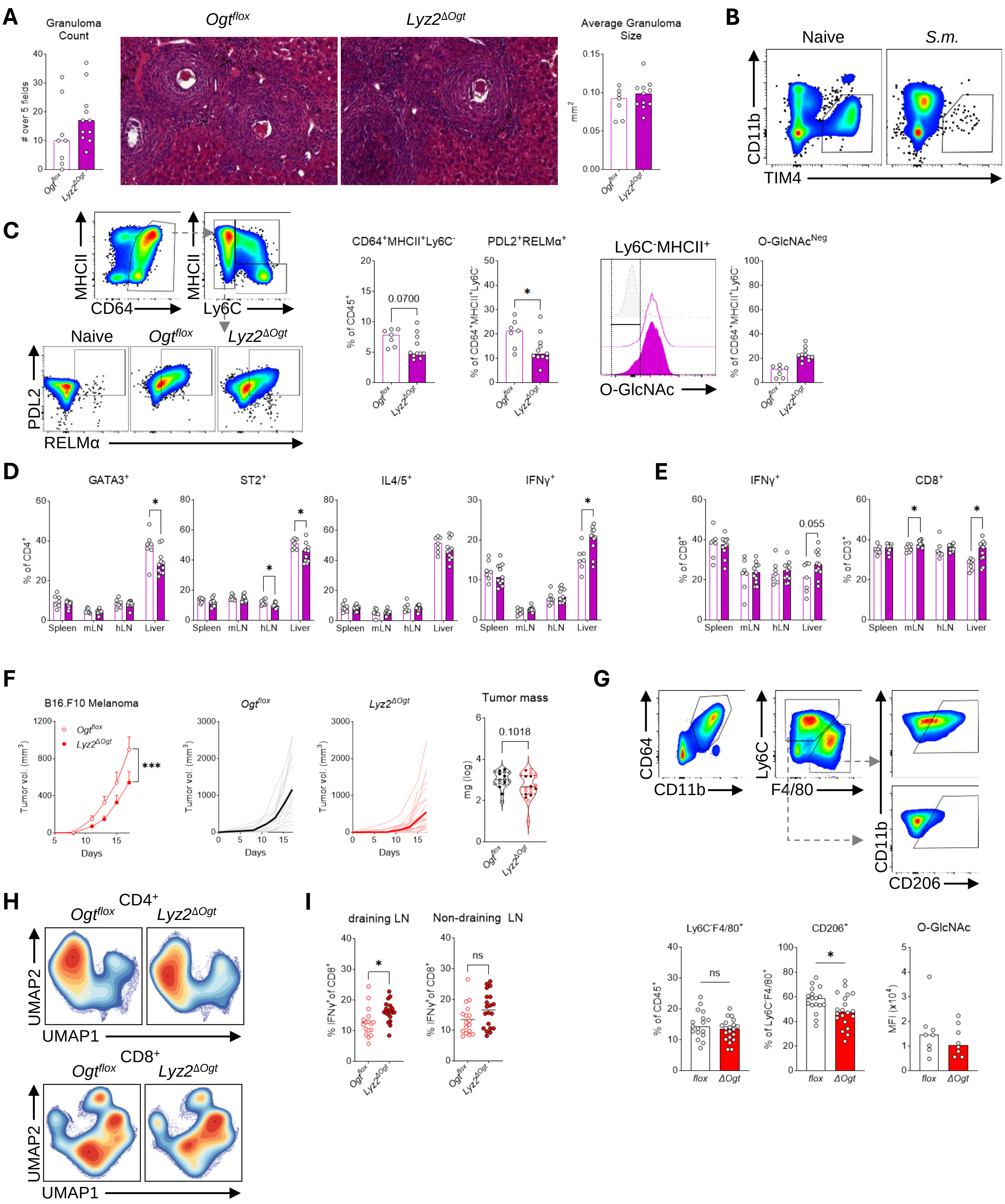

### Figure S3 – related to main Figure 3

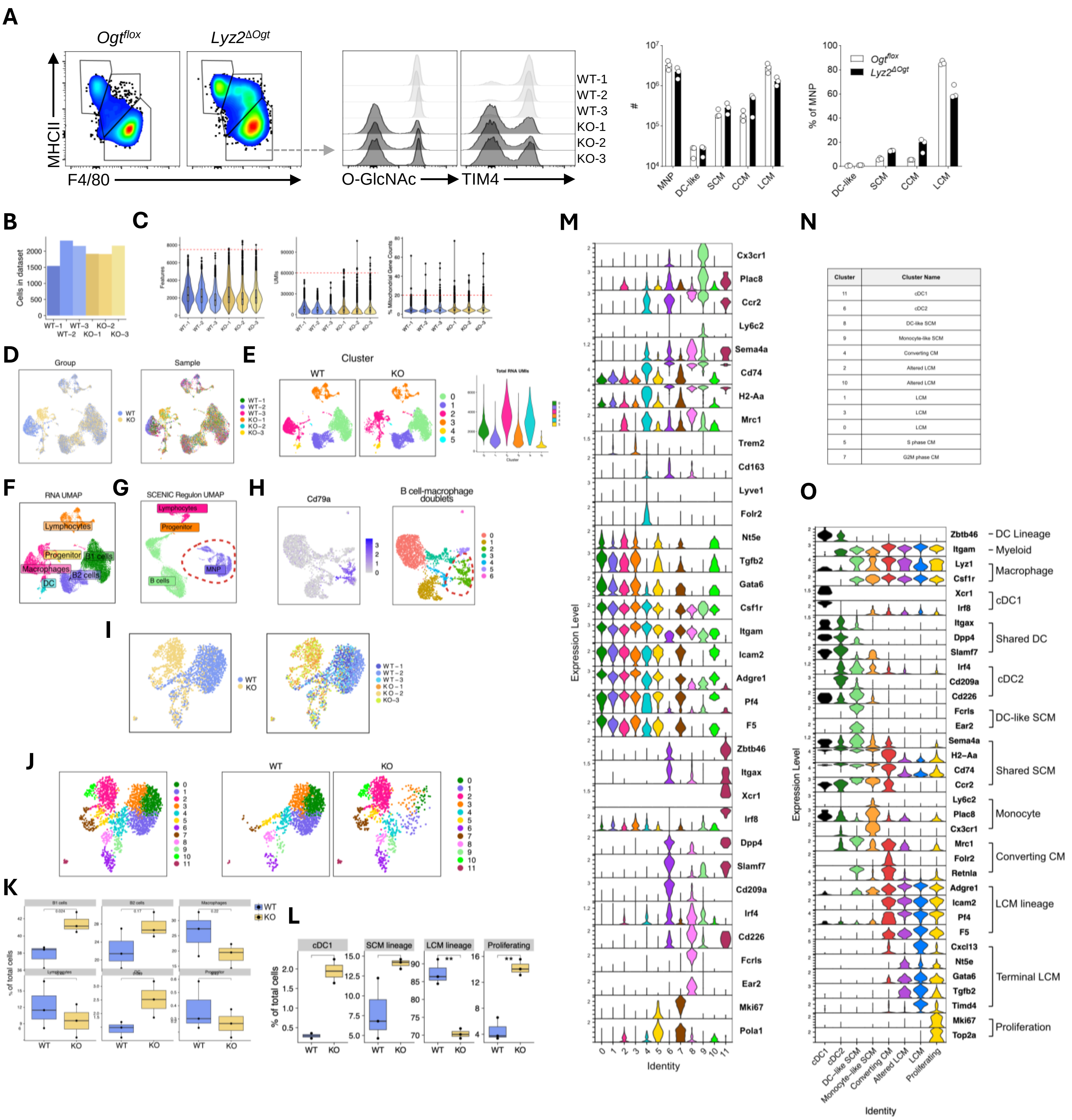


| Cluster | Cluster Name |
|---|---|
| 11 | cDC1 |
| 6 | cDC2 |
| 8 | DC-like SCM |
| 9 | Monocyte-like SCM |
| 4 | Converting CM |
| 2 | Altered LCM |
| 10 | Altered LCM |
| 1 | LCM |
| 3 | LCM |
| 0 | LCM |
| 5 | S phase CM |
| 7 | G2M phase CM |

### Figure S4 – Related to main Figure 4

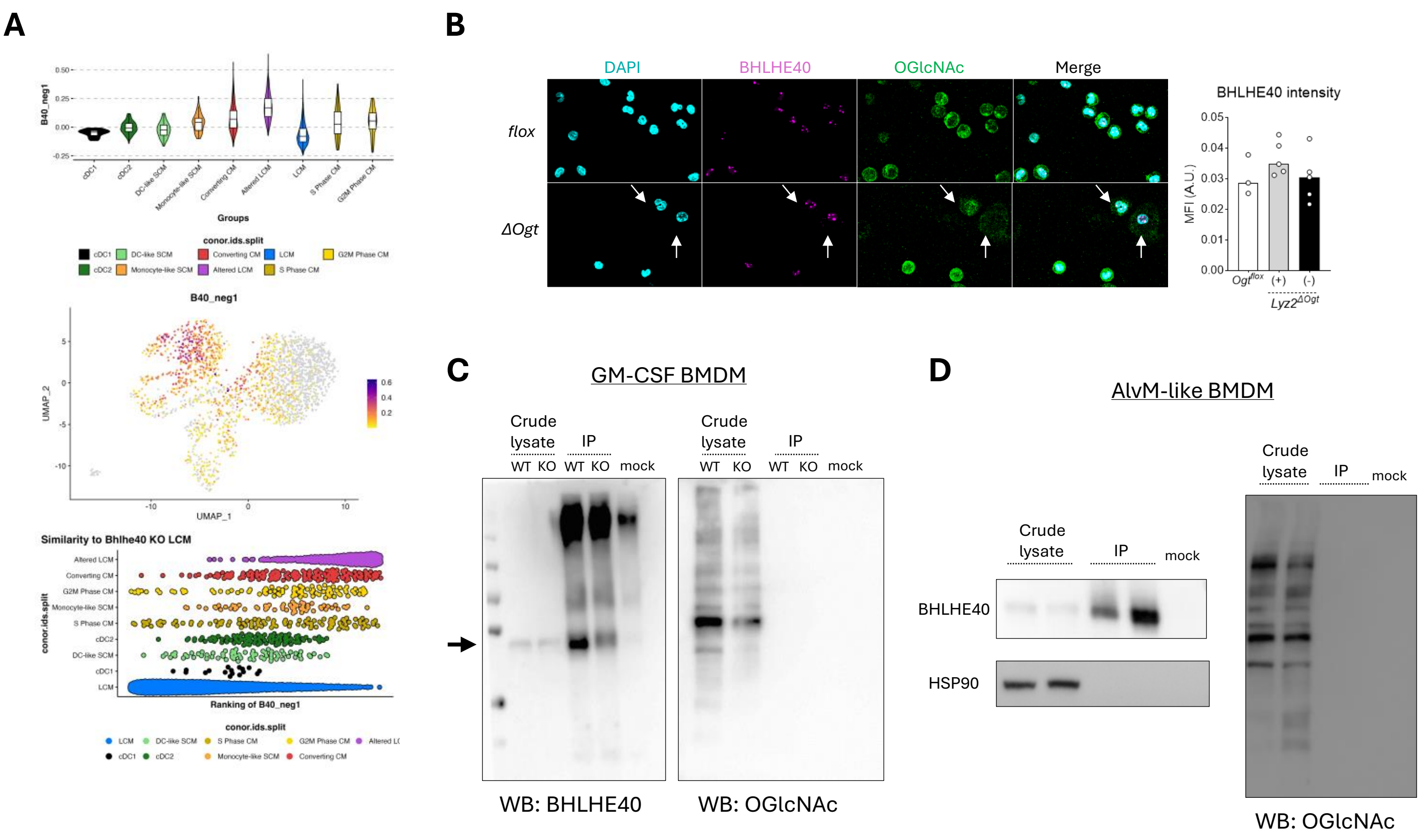

### Figure S5 – Related to main Figures 5 & 6

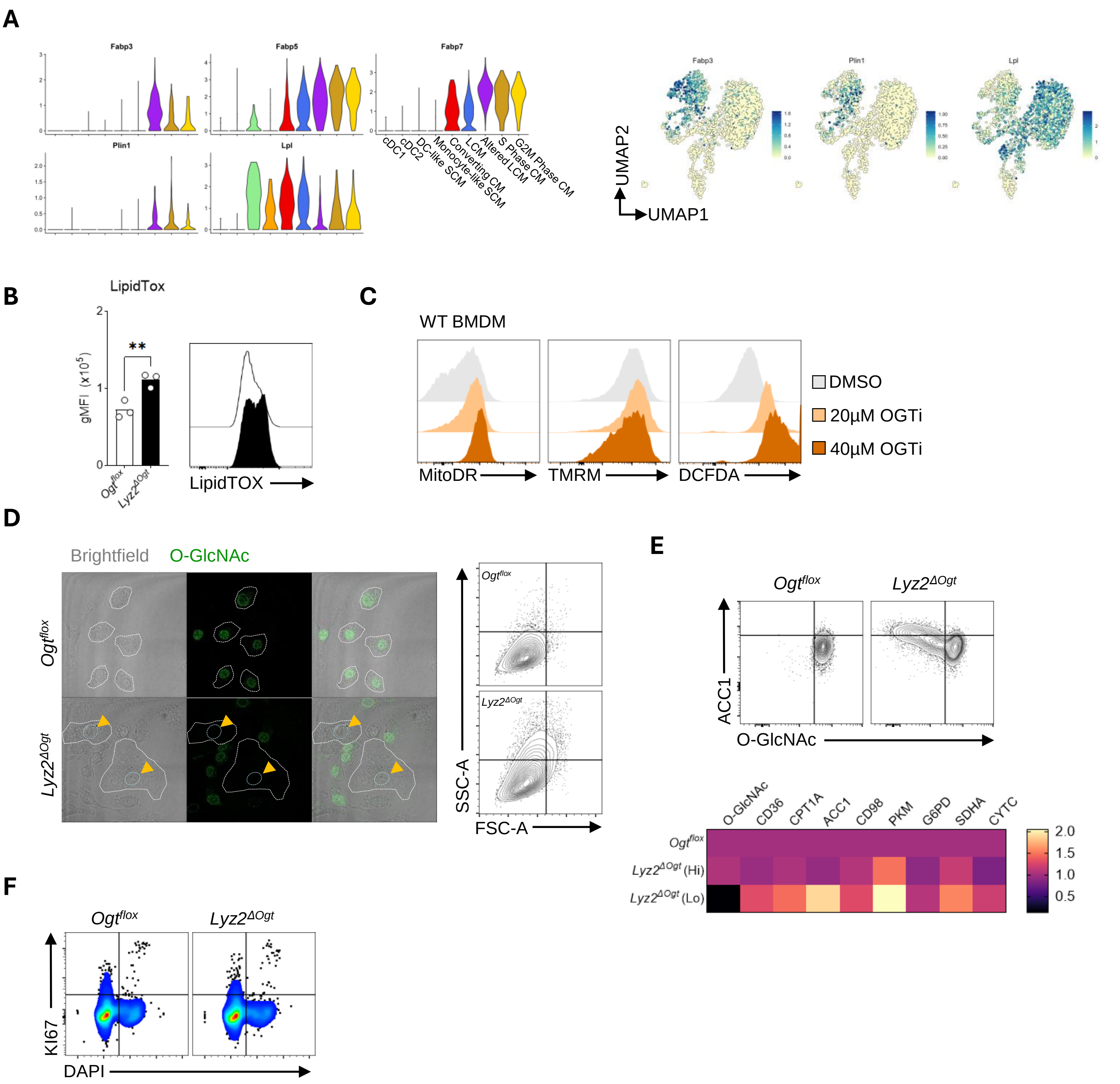

### Figure S6 – Related to main Figure 7

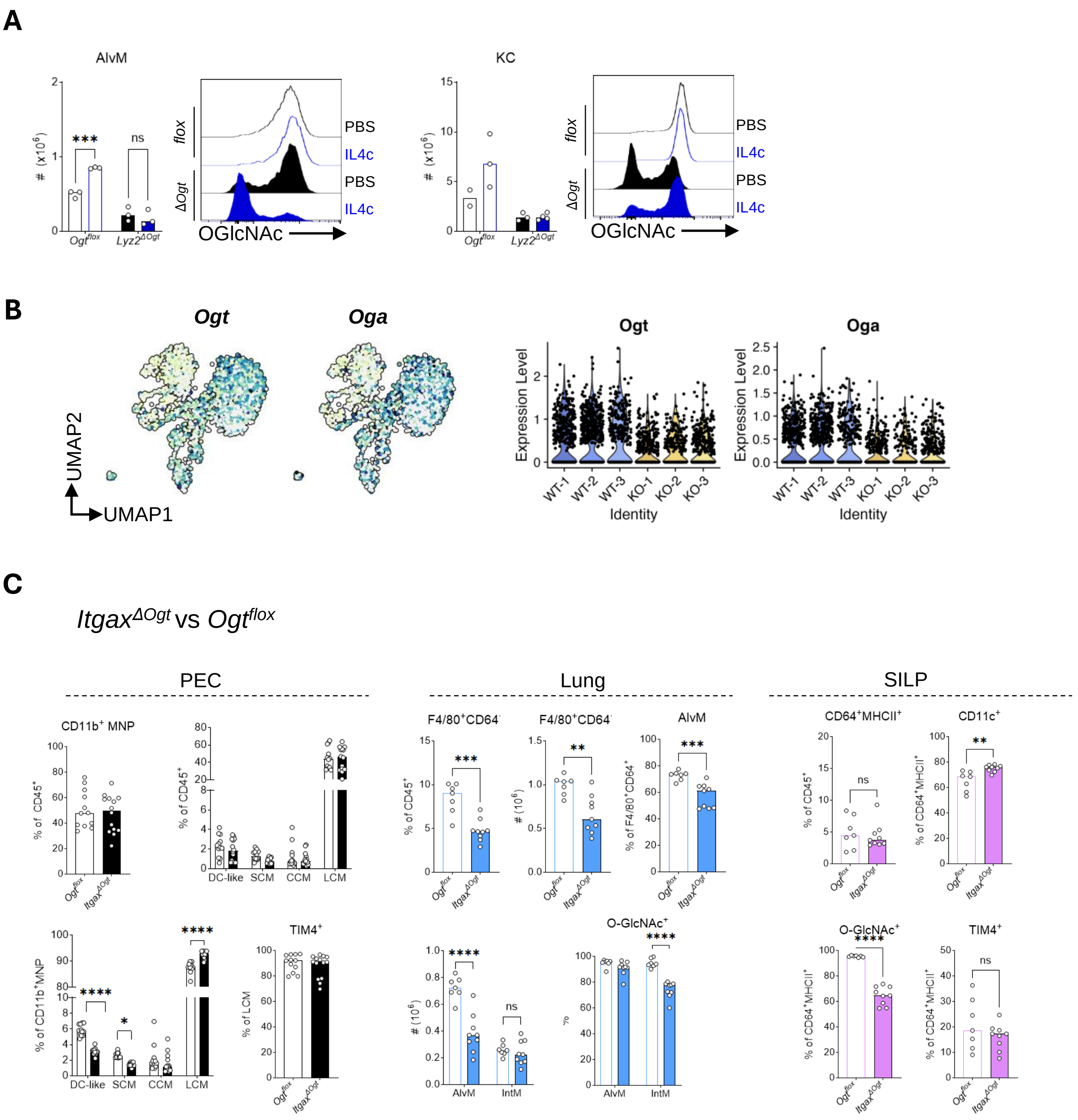

### Figure S7 – Related to main Figure 7

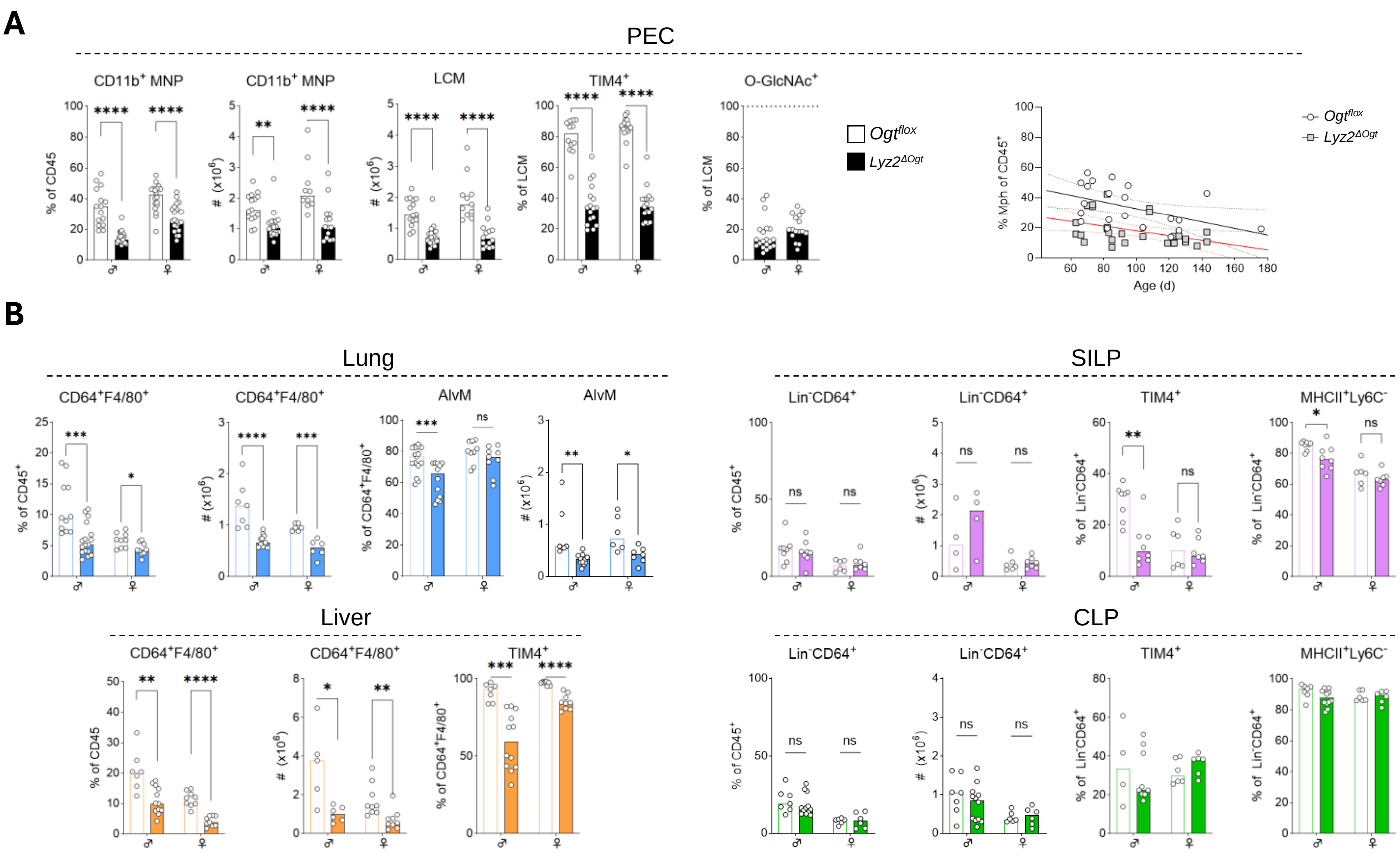
